# The mitotic surveillance pathway requires PLK1-dependent 53BP1 displacement from kinetochores

**DOI:** 10.1101/2023.03.27.534346

**Authors:** Matteo Burigotto, Vincenza Vigorito, Alessia Mattivi, Colin Gliech, Sabrina Ghetti, Alessandra Bisio, Graziano Lolli, Andrew J. Holland, Luca L. Fava

## Abstract

53BP1 acts at the crossroads between DNA repair and p53-mediated stress response. With its interactor USP28, it is part of the mitotic surveillance pathway (MSP), a sensor that monitors the duration of cell division, promoting p53-dependent cell cycle arrest when a critical time threshold is surpassed. 53BP1 dynamically associates with kinetochores, being recruited during prophase, and then undergoing a time-dependent loss of affinity. However, the relevance of this behaviour remains unclear. Here, we identify CENP-F as an interaction partner and kinetochore receptor for 53BP1. By engineering human cells with a CENP-F point mutation, we demonstrate that preventing 53BP1 kinetochore localization does not reduce MSP proficiency. Strikingly, however, preventing the loss of 53BP1 from the kinetochore by inhibiting Polo-like kinase 1 (PLK1) restrains MSP activity, a phenomenon that is abrogated in the CENP-F mutant condition. Taken together, we demonstrate that kinetochore-loaded 53BP1 represents an MSP functionally inhibited state and that PLK1-dependent re-localization of 53BP1 represents an important layer of MSP regulation.

## Introduction

Development and homeostasis of multicellular organisms critically depend on the fidelity of cell division, granting genome stability over a multitude of subsequent mitoses. Mitosis is a highly coordinated set of events involving the abrupt functional and morphological reorganization of the cell in a short time, ultimately leading to the generation of two genetically identical cells. Mitotic duration is determined by the lag between Cyclin B/CDK1 activation, kick-starting mitotic entry, and the complete activation of an E3 ubiquitin ligase called the anaphase-promoting complex/cyclosome (APC/C), which in turn promotes mitotic exit. Staggered waves of Cyclin B/CDK1 and APC/C activity constitute the core cell cycle clock, driving the alternance between interphase and mitosis (Murray & Kirschner, 1989; Pines, 2011). However, the timing of cell division also depends on an additional rheostat: the spindle assembly checkpoint (SAC). The SAC acts by delaying APC/C activation until all chromosomes are bi-oriented on the mitotic spindle, thereby lowering the frequency of chromosome segregation errors (Collin *et al*, 2013).

Due to the high metabolic demand of the mitotic status and the inability of the SAC to completely hinder Cyclin B degradation, an arrest in mitosis cannot be sustained indefinitely (Doménech *et al*, 2015; Brito & Rieder, 2006). Moreover, merotelic kinetochore-microtubule attachments cannot always be corrected by the activity of the SAC (Gregan *et al*, 2011; Dudka *et al*, 2018). Thus, damaged or stressed cells often undergo an extension of the mitotic duration that is mediated by the SAC, yet this might not suffice to grant complete fidelity. Seminal work by Uetake and Sluder revealed the presence of an additional fail-safe mechanism by demonstrating that mitoses whose duration exceeds a critical threshold (90 minutes in their experimental conditions) yield a progeny that become arrested in the subsequent G1 phase in a p53-dependent manner (Uetake & Sluder, 2010). Moreover, three independent genetic screens revealed that the activation of the p53-p21 axis upon centrosome depletion relies on two additional factors, namely the oligomeric multidomain scaffold 53BP1 and the ubiquitin-specific protease USP28 (Fong *et al*, 2016; Lambrus *et al*, 2016; Meitinger *et al*, 2016). 53BP1 and USP28 display the ability to form a ternary protein complex with p53 and support its ability to transactivate target genes such as p21 (Cuella-Martin *et al*, 2016). This pathway is commonly called the mitotic surveillance pathway (MSP). The MSP is operational in somatic cell divisions and has measurable activity during embryonic development starting at around E7 in mice (Xiao *et al*, 2021). Moreover, in mouse models of microcephaly, in which mitotic delay in neural progenitor cells is achieved by deletion of centrosomal proteins, aberrant activation of the MSP appeared crucial to the aetiology of the microcephalic phenotype (Phan *et al*, 2021; Phan & Holland, 2021). However, the mechanism governing MSP complex assembly and activity remains unclear.

Kinetochores (KTs) are multi-protein assemblies built on centromeric loci, pivotal for the interactions between sister chromatids and the spindle microtubules (Pesenti *et al*, 2016). In addition to merely structural roles, they are important for SAC signalling, thereby defining the mitotic duration (Foley & Kapoor, 2013; Lara-Gonzalez *et al*, 2021). Strikingly, 53BP1 was reported to localize to the outermost layer of KTs, known as the fibrous corona, the same substructure responsible for igniting the SAC (Jullien *et al*, 2002; Fong *et al*, 2016; Lambrus *et al*, 2016). Moreover, 53BP1 association with KTs appears to be dependent on the duration of mitosis, but is independent on the activation status of the SAC (Fong *et al*, 2016). Whether the transient association of 53BP1 with KTs or its subsequent dissociation are important for MSP activation remains to be established. Here, we demonstrate that the fibrous corona protein CENP-F and 53BP1 interact directly and that this is key to 53BP1 recruitment at the KT. Using gene editing, we introduced a single amino acidi substitution into the endogenous *CENPF* locus in human cells. Exploiting this mutant, we show that i) KT-localization of 53BP1 is not necessary for MSP functionality, and ii) sequestering 53BP1 at KTs by interfering with PLK1 activity prevents MSP activation. Thus, we conclude that although not sufficient to fully activate the MSP, 53BP1 release from KTs is an essential step in MSP activation.

## Results & Discussion

### CENP-F is the 53BP1 KT receptor

To unbiasedly characterize the KT receptor of 53BP1, we employed CRISPR/Cas9 (Ghetti *et al*, 2021) to engineer the endogenous *TP53BP1* locus of hTERT-RPE1 cells by introducing a biallelic V5-epitope tag at the C-terminus of the protein (Fig. 1A). V5-tagged 53BP1 displayed a behaviour indistinguishable from the untagged protein, showing a pan-nuclear distribution with some discrete foci in interphase cells, and being recruited at KTs upon mitotic entry (Fig. 1B). Moreover, the 53BP1-V5 fusion protein mirrored the untagged protein, accumulating at γH2AX-positive foci in irradiated cells (Fig. EV1A-B and Appendix Fig. A-D) and displaying similar mitotic phase-dependent dynamics at the KT, peaking in prophase and gradually disappearing from this structure during mitotic progression (Fig. EV1C). Thus, the tagging of 53BP1 with V5 did not interfere with protein-protein interactions relevant to its cell cycle dynamic localization.

**Figure 1.**
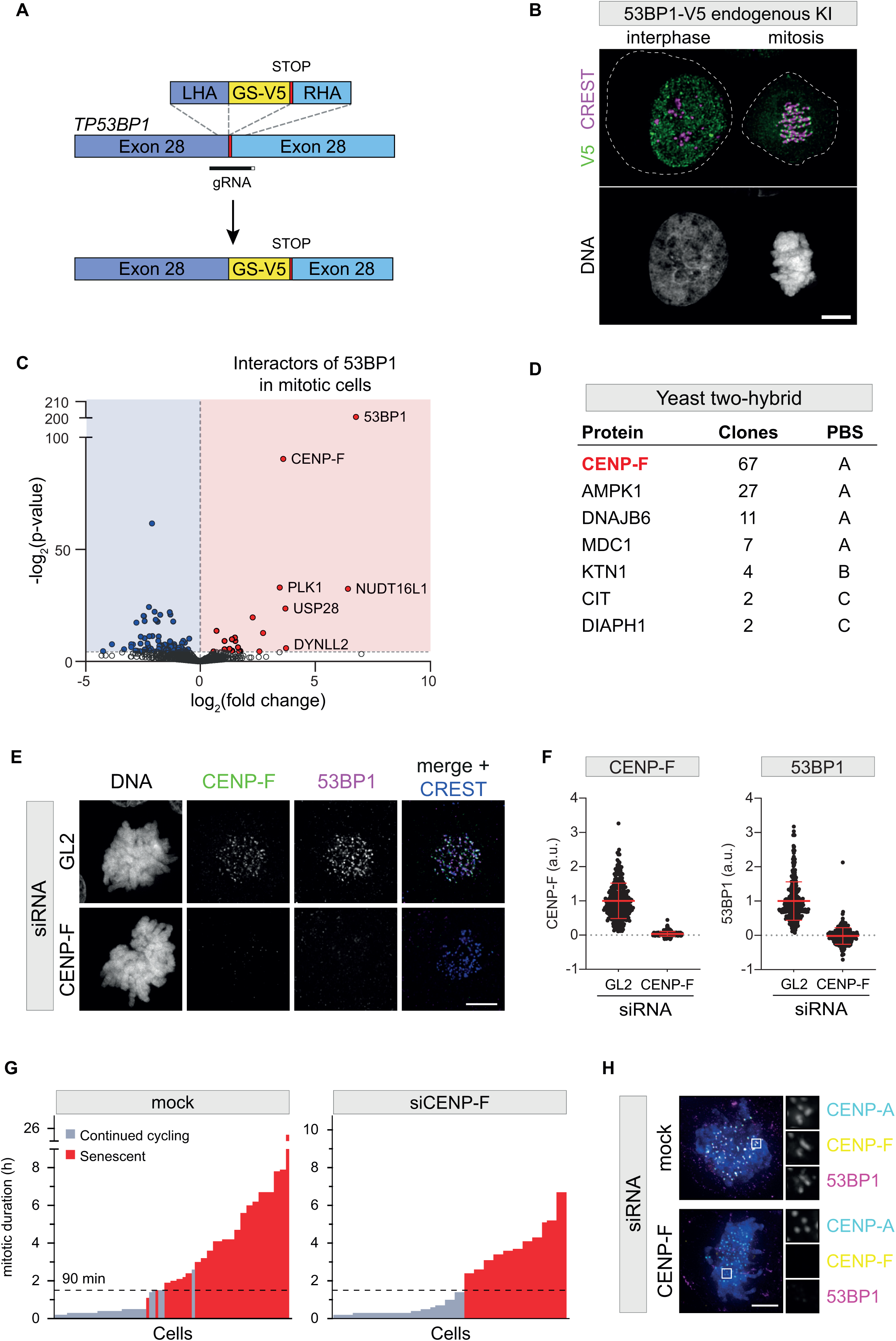
CENP-F is the 53BP1 kinetochore receptor. A) Schematic of the knock-in strategy to introduce the V5 sequence into the endogenous *TP53BP1* locus. LHA: left homology arm; RHA: right homology arm; GS-V5: Gly-Ser linker followed by V5-epitope tag. B) Representative fluorescence micrograph of 53BP1-V5 cells co-stained with the indicated antibodies. An interphase (left) and a mitotic cell (right) are shown. The dashed lines indicate the plasma membrane of the two cells. Scale bar: 5 µm. C) Volcano plot showing mitosis-specific 53BP1 interacting proteins identified by mass spectrometry. Each dot depicts a single protein. The x axis represents the log_2_(fold change) over the untagged control; the y axis shows significance expressed as -log_10_(p-value). D) List of the main prey fragments retrieved by the yeast two-hybrid screen. The name of the protein, the number of clones, along with the specific protein binding score (PBS) are reported. E) HeLa S3 cells were transfected with the indicated siRNA and subjected to immunofluorescence using the indicated antibodies. Scale bar: 5 µm. F) Dot plots showing the intensity of the indicated proteins at individual KTs. Mean values (red lines) ± SD are reported. Data obtained from images as in E). N ≥ 295 KTs were assessed from at least 20 cells for each condition; a.u. = arbitrary units. G) Senescence threshold plots of RPE1 p21-EGFP H2B-iRFP cells transiently treated with dimethylenastron after CENP-F or mock siRNA. 77 control cells and 48 CENP-F depleted cells were imaged for 2 days. H) RPE1 p21-EGFP H2B-iRFP cells were transfected with the indicated siRNA and subjected to immunofluorescence using the indicated antibodies. Blow-ups are magnified 3.5X. Scale bar: 5 µm.

To define the mitotic interactors of 53BP1, we synchronized RPE1 53BP1-V5 knock-in cells in prometaphase and performed immunoprecipitation against the V5-tag, followed by LC-MS-mediated identification of the proteins specifically enriched in comparison to immunoprecipitation performed against RPE1 parental cells treated in the same fashion. MS analysis detected 21 specific binding partners (Table 1), including some known 53BP1 interactors, such as NUDT16L1/TIRR, DYNLL2, USP28, and PLK1 (Fig. 1C). The KT fibrous corona protein CENP-F scored among the top enriched hits (Fig. 1C), suggesting that it may play a role in the localization of 53BP1 at the KT. In an orthogonal search to define the KT-binding partner(s) of 53BP1, we performed a yeast two-hybrid screen using the KT-binding domain of 53BP1 (Jullien *et al*, 2002) as bait against a human cDNA library. Of the 173 processed clones, 67 CENP-F clones were identified at a very high confidence (Protein Binding Score of A) (Formstecher *et al*, 2005), thereby confirming that CENP-F and 53BP1 are direct *bona fide* interactors (Fig. 1D).

**Table 1.**
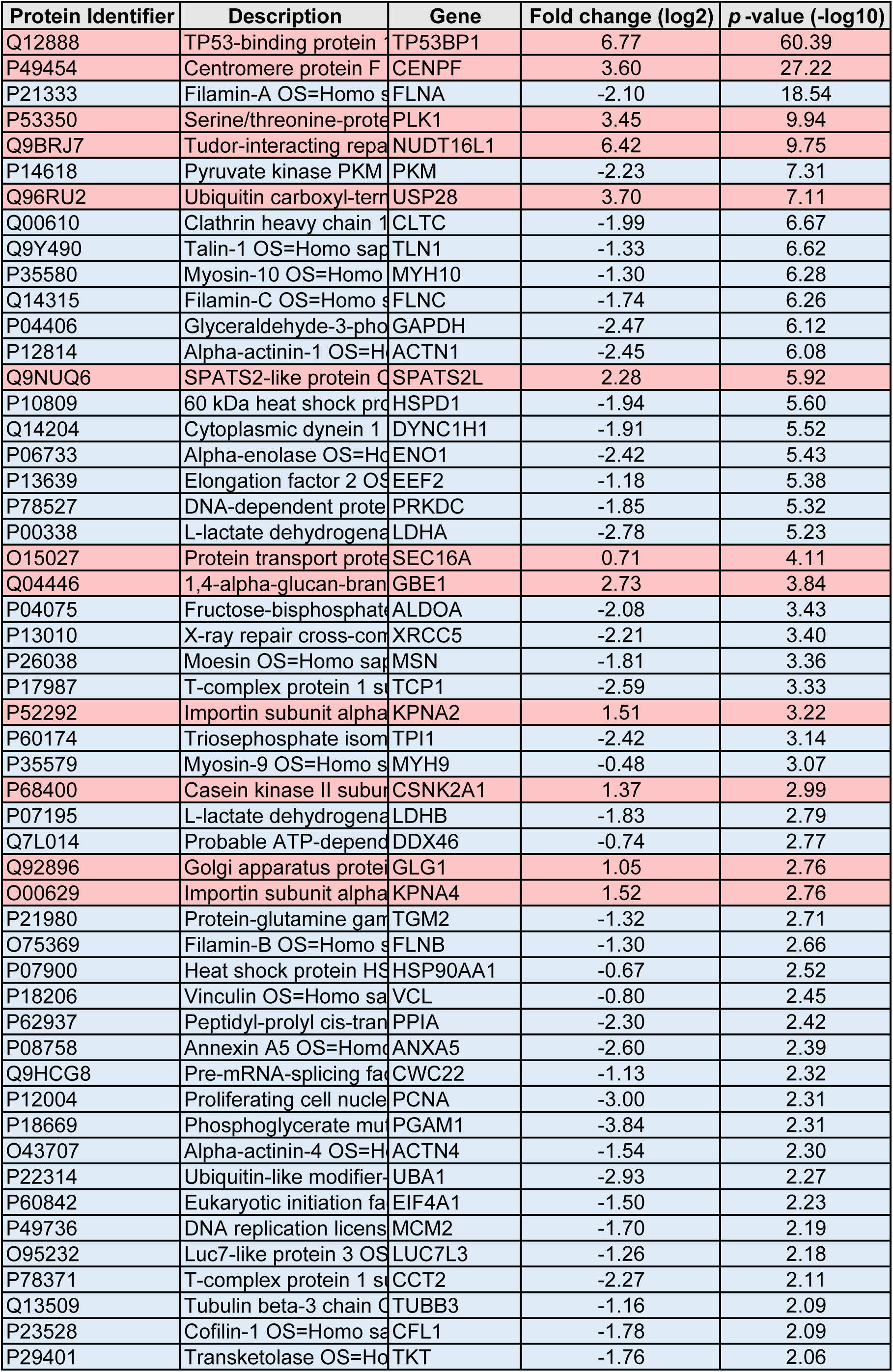
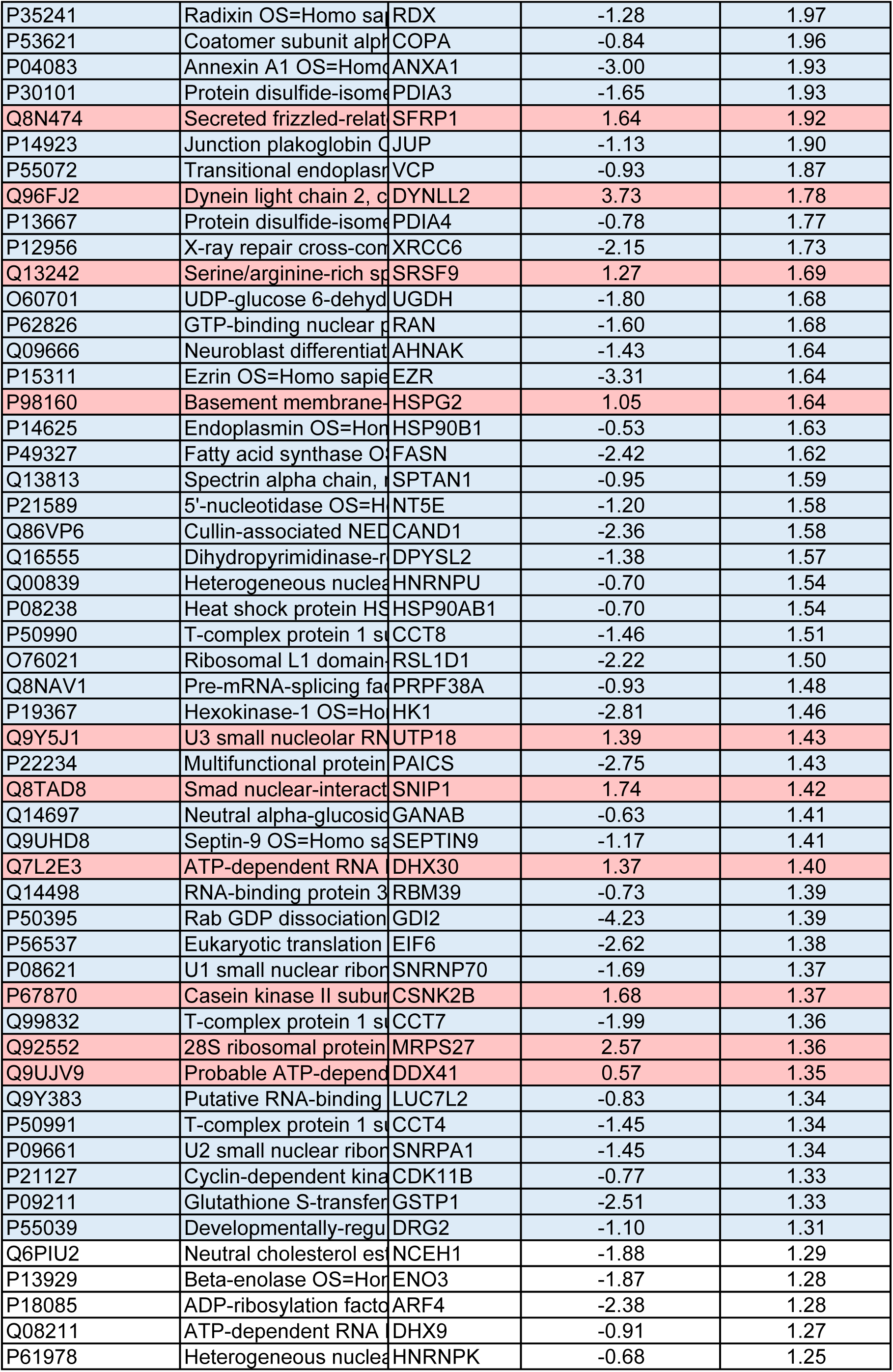
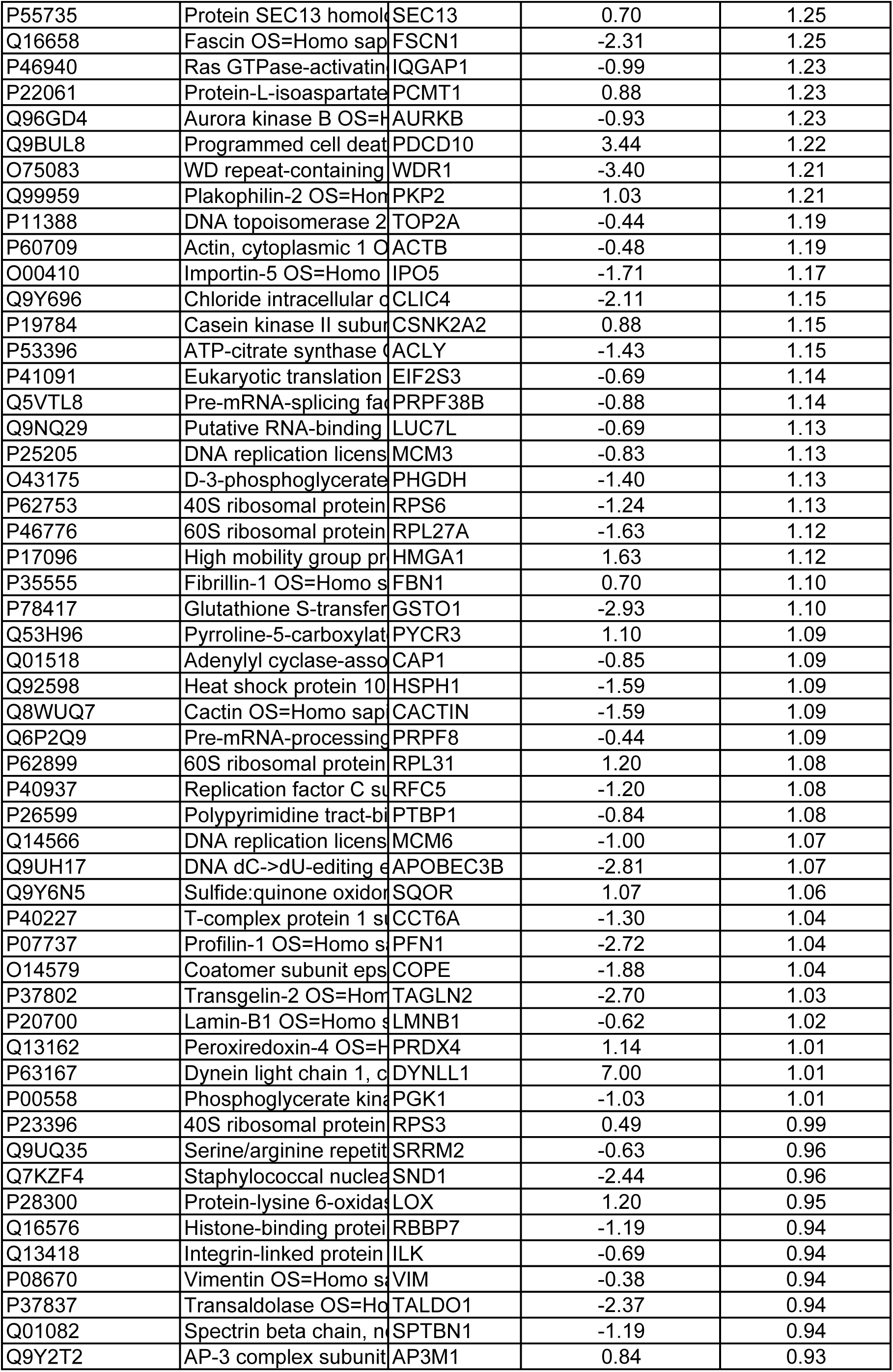
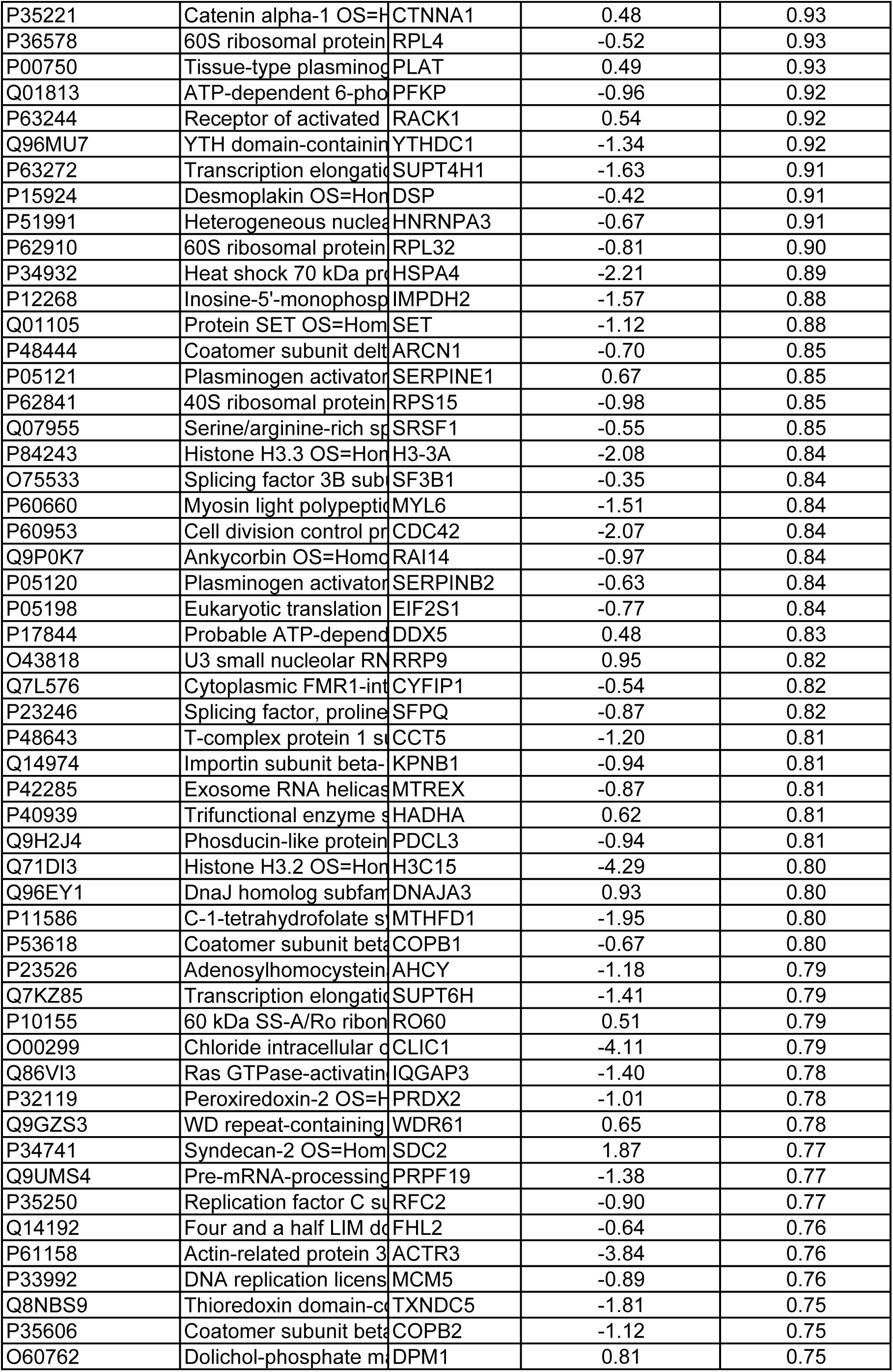
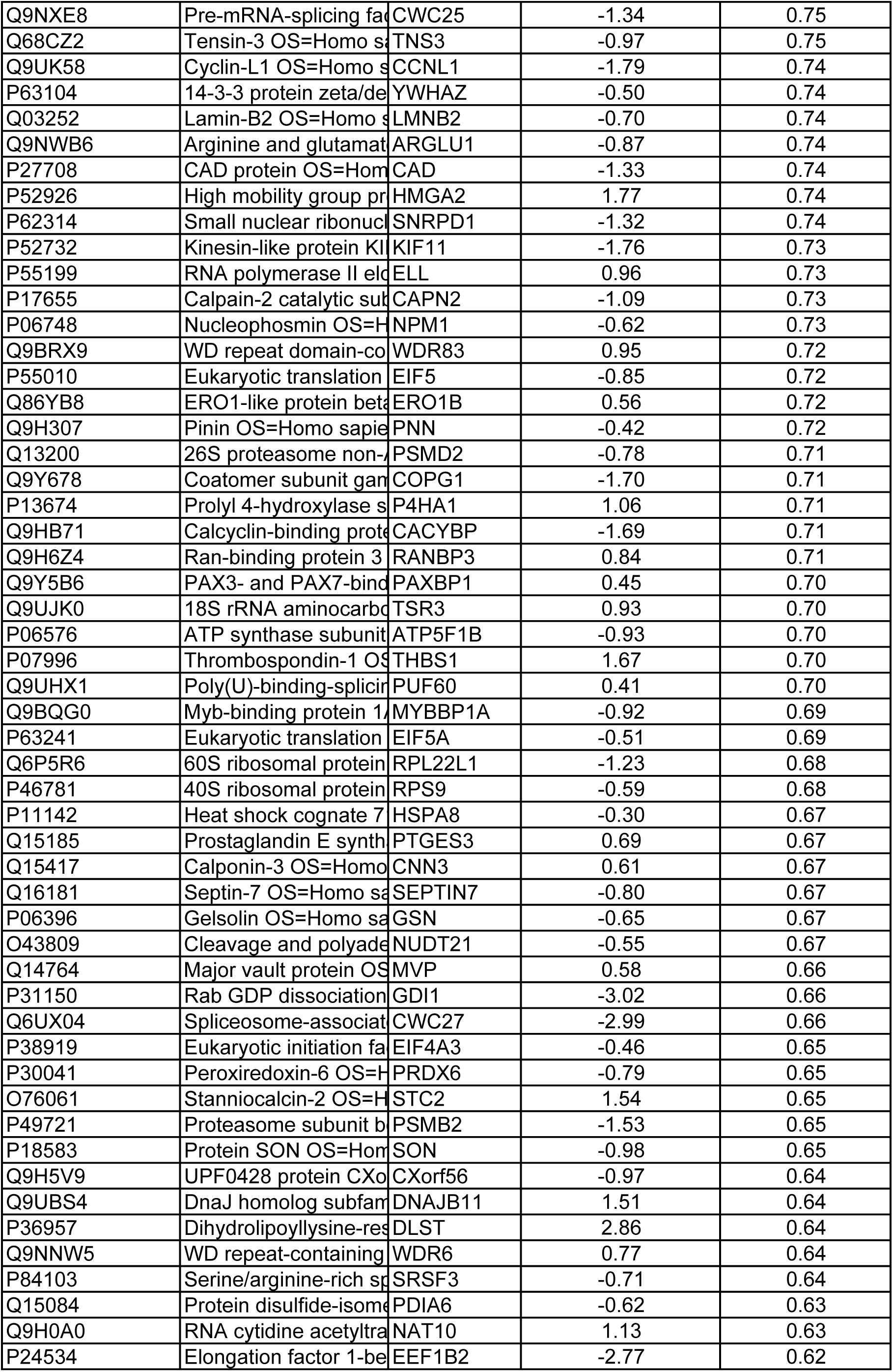
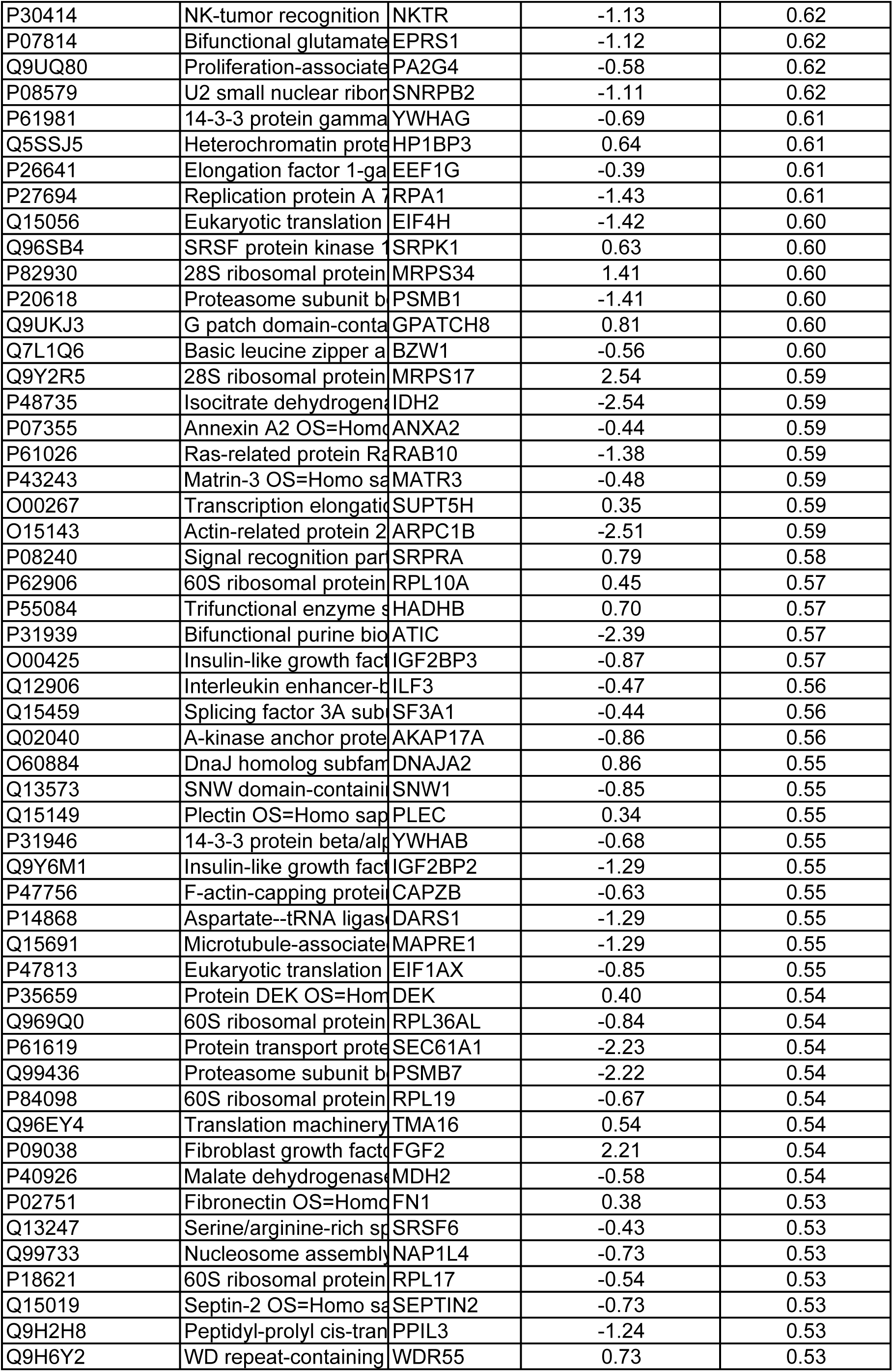
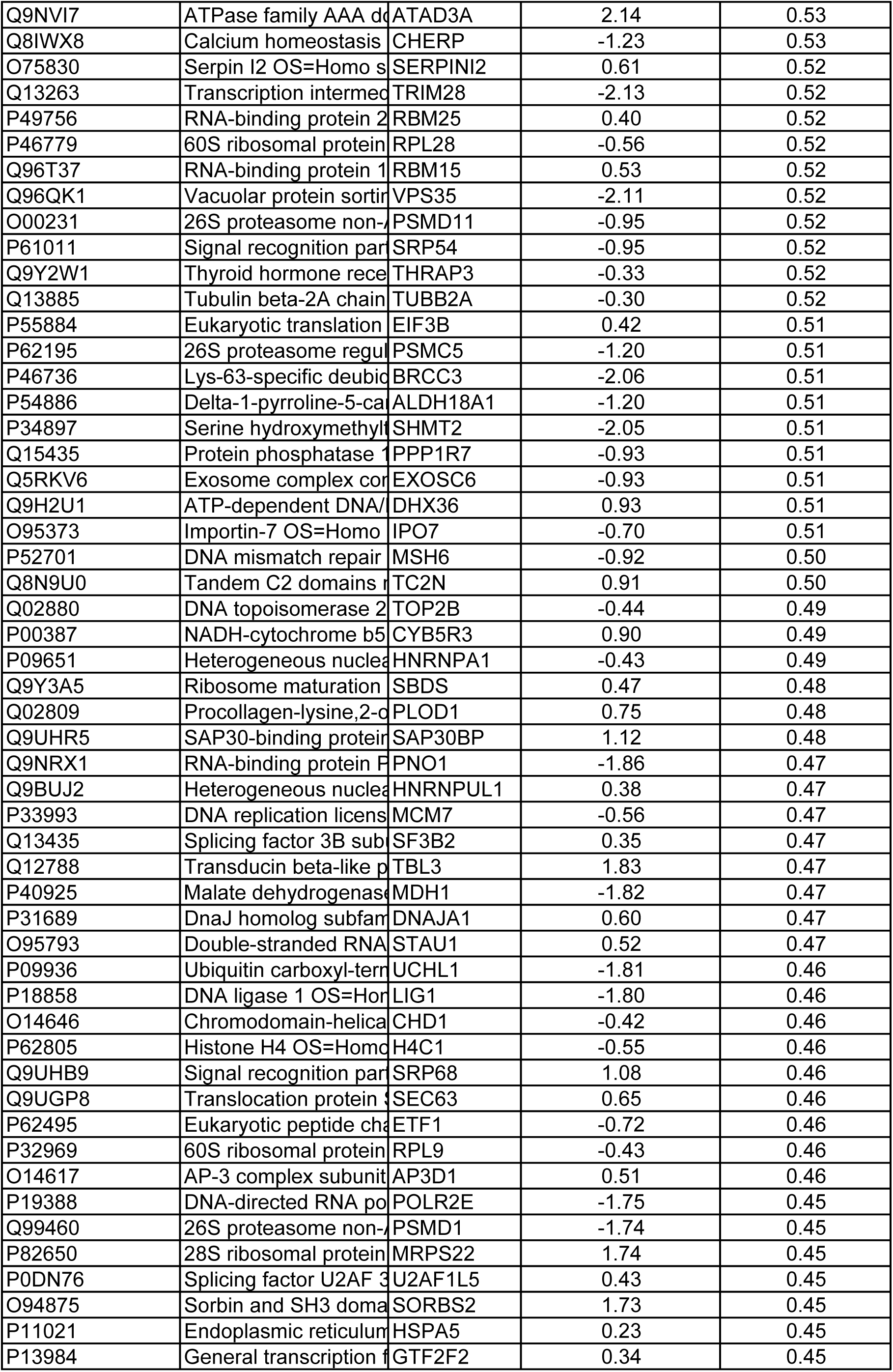
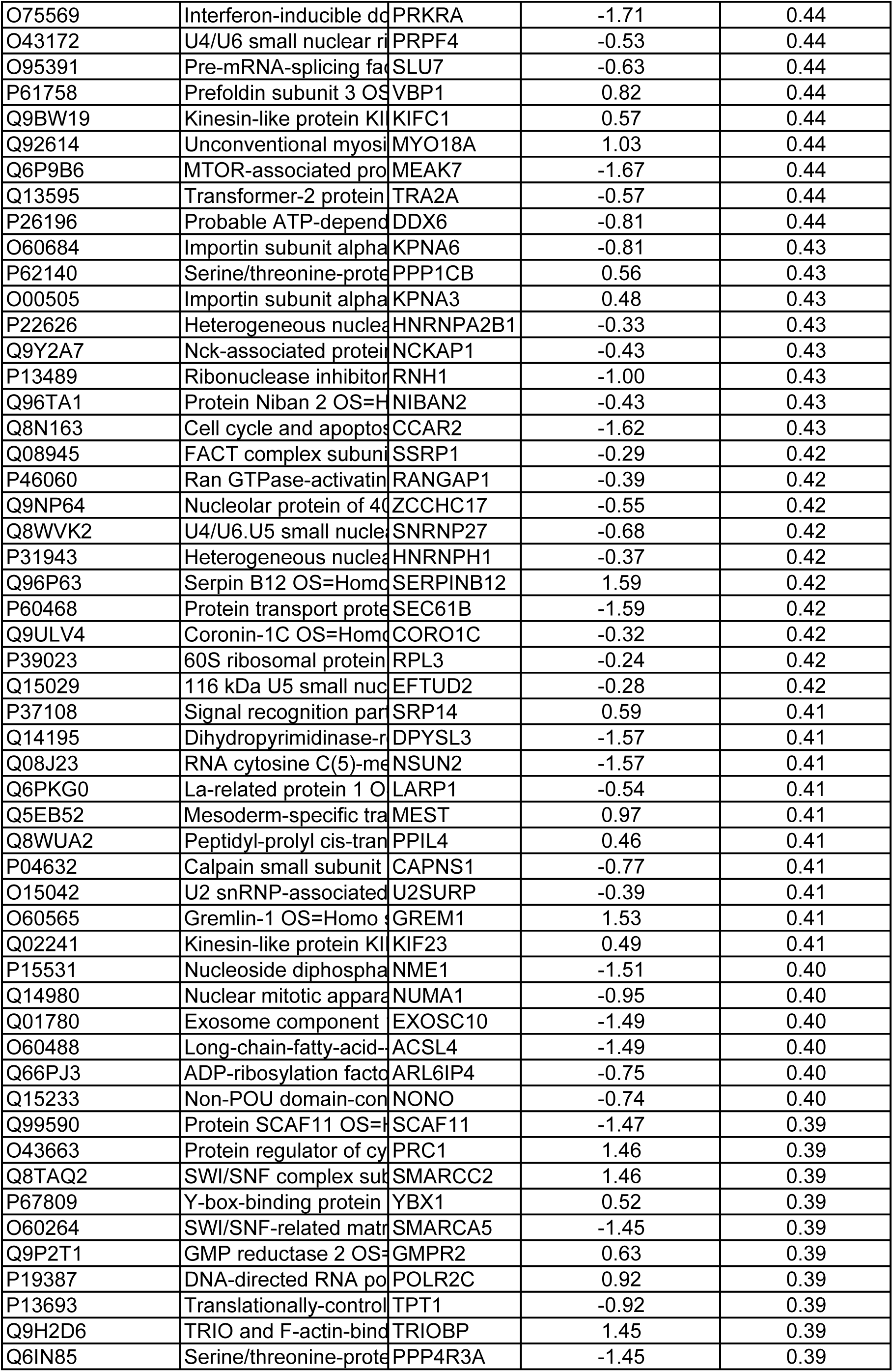
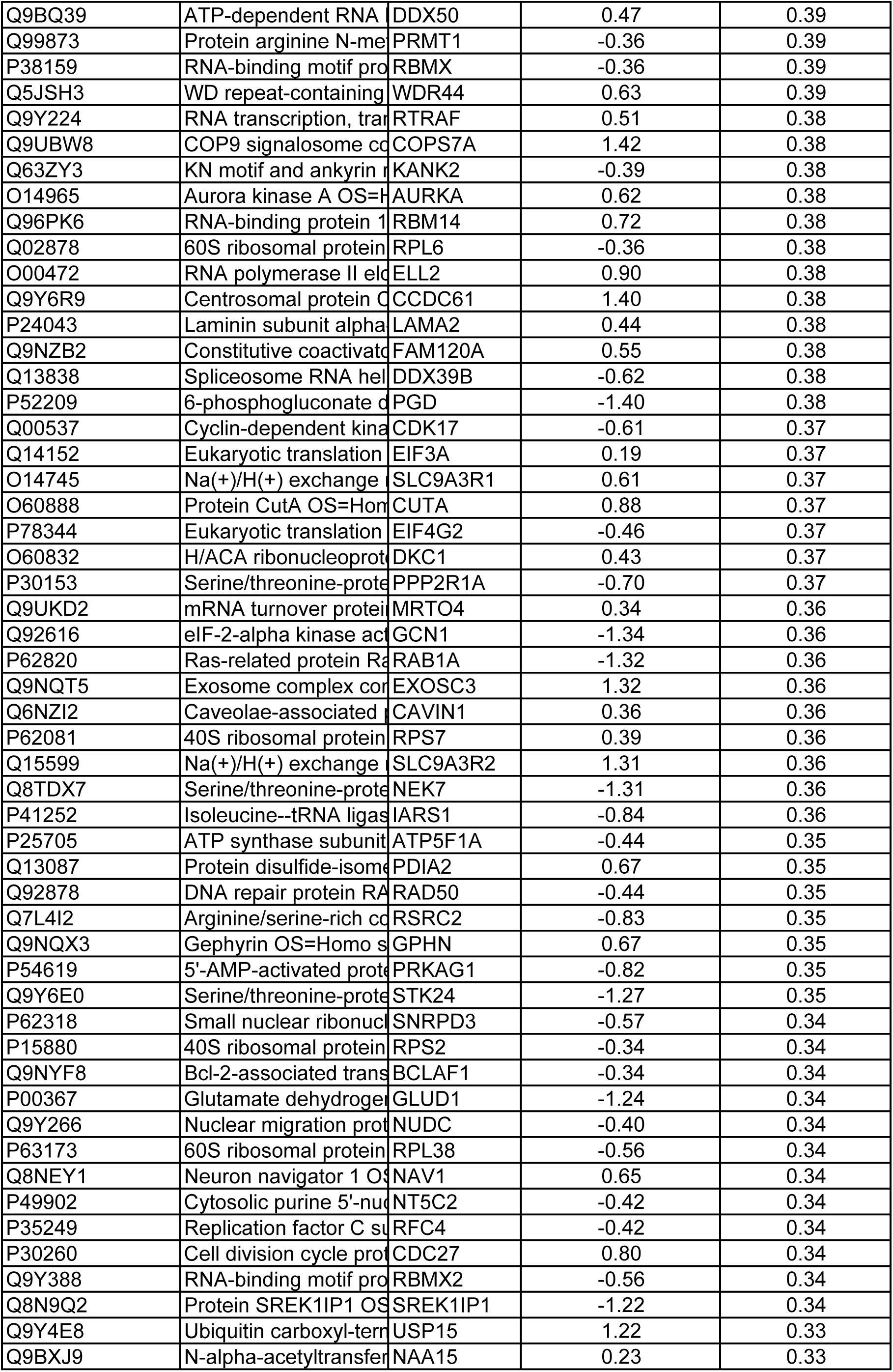
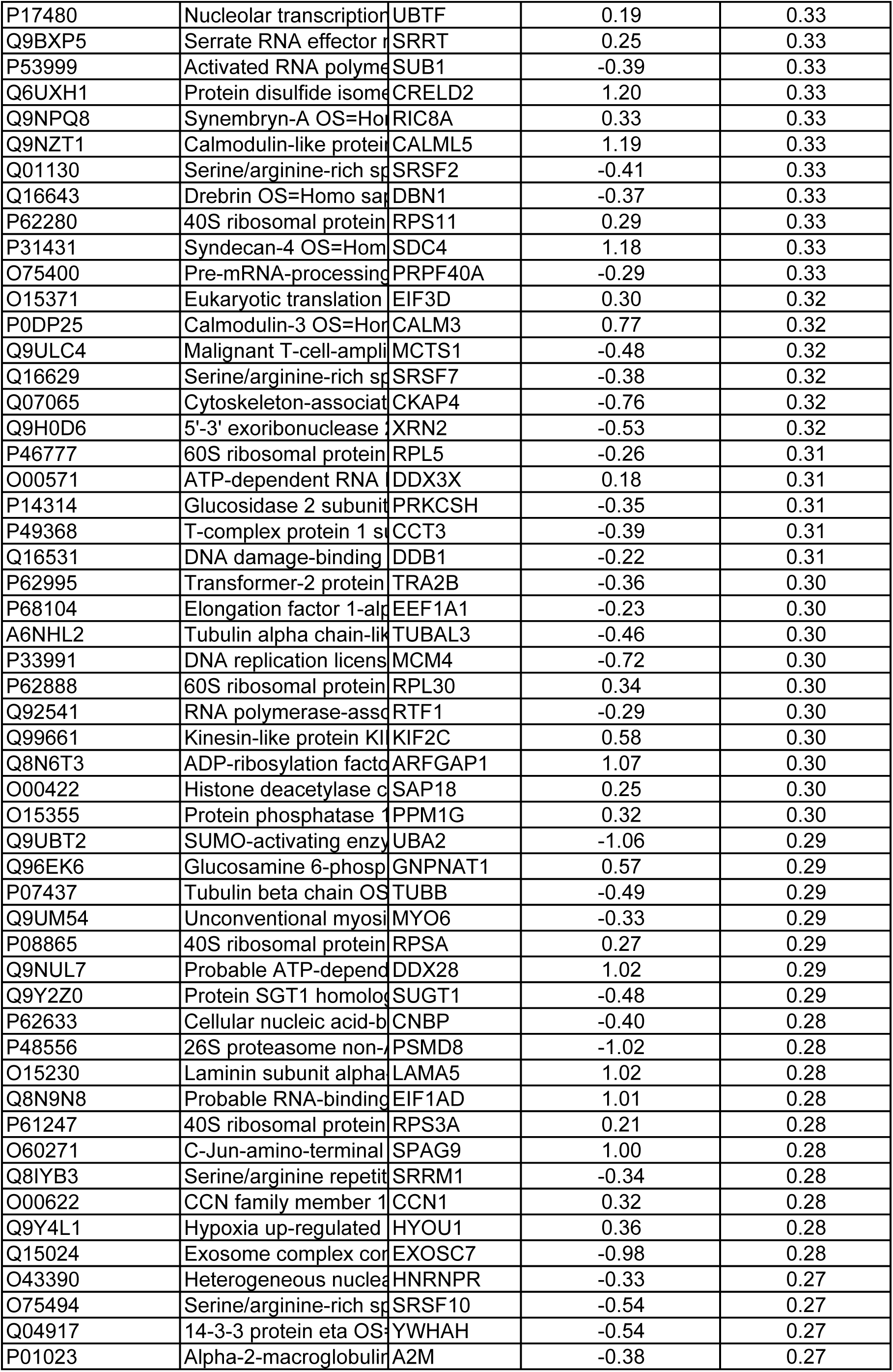
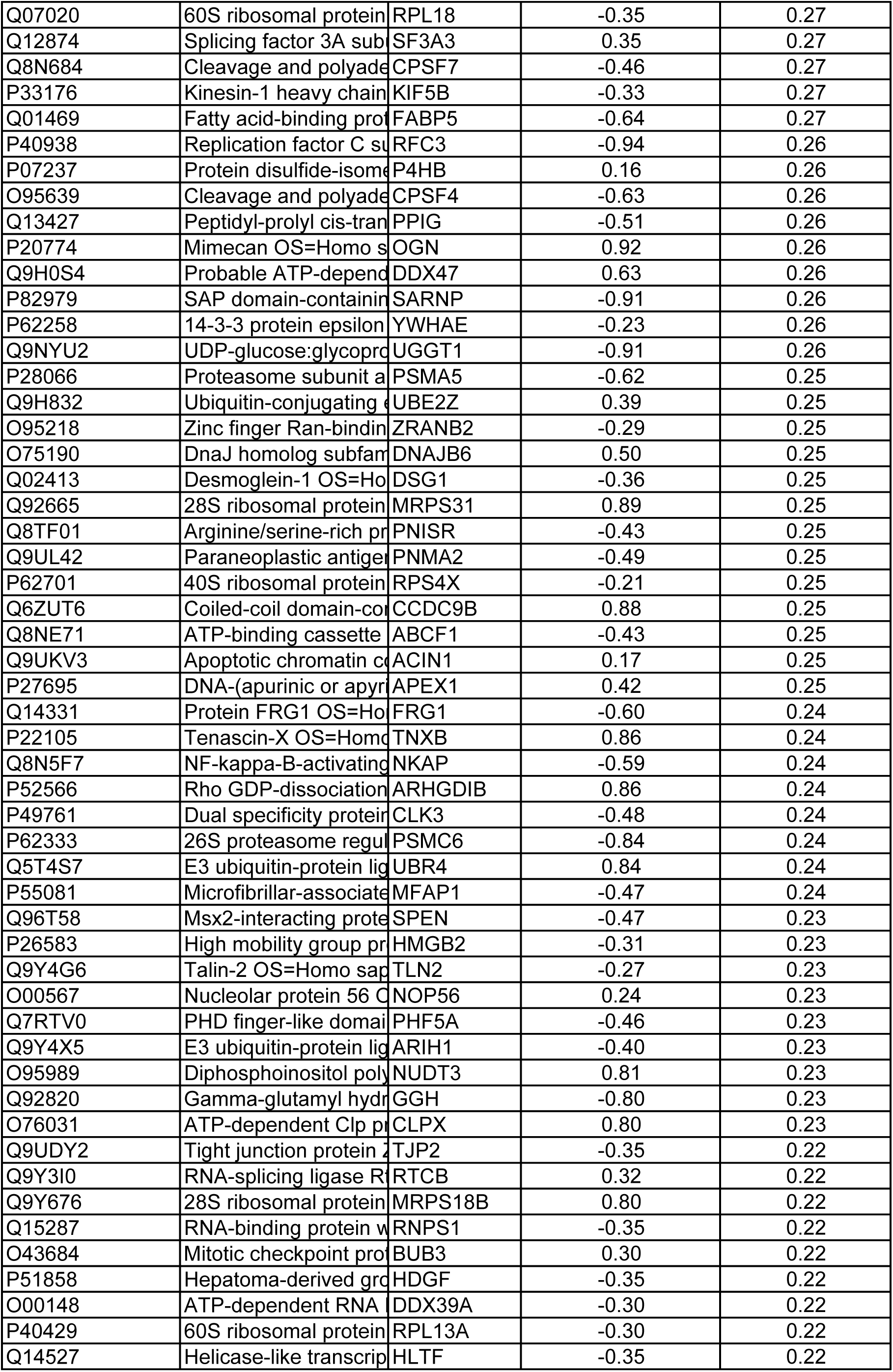
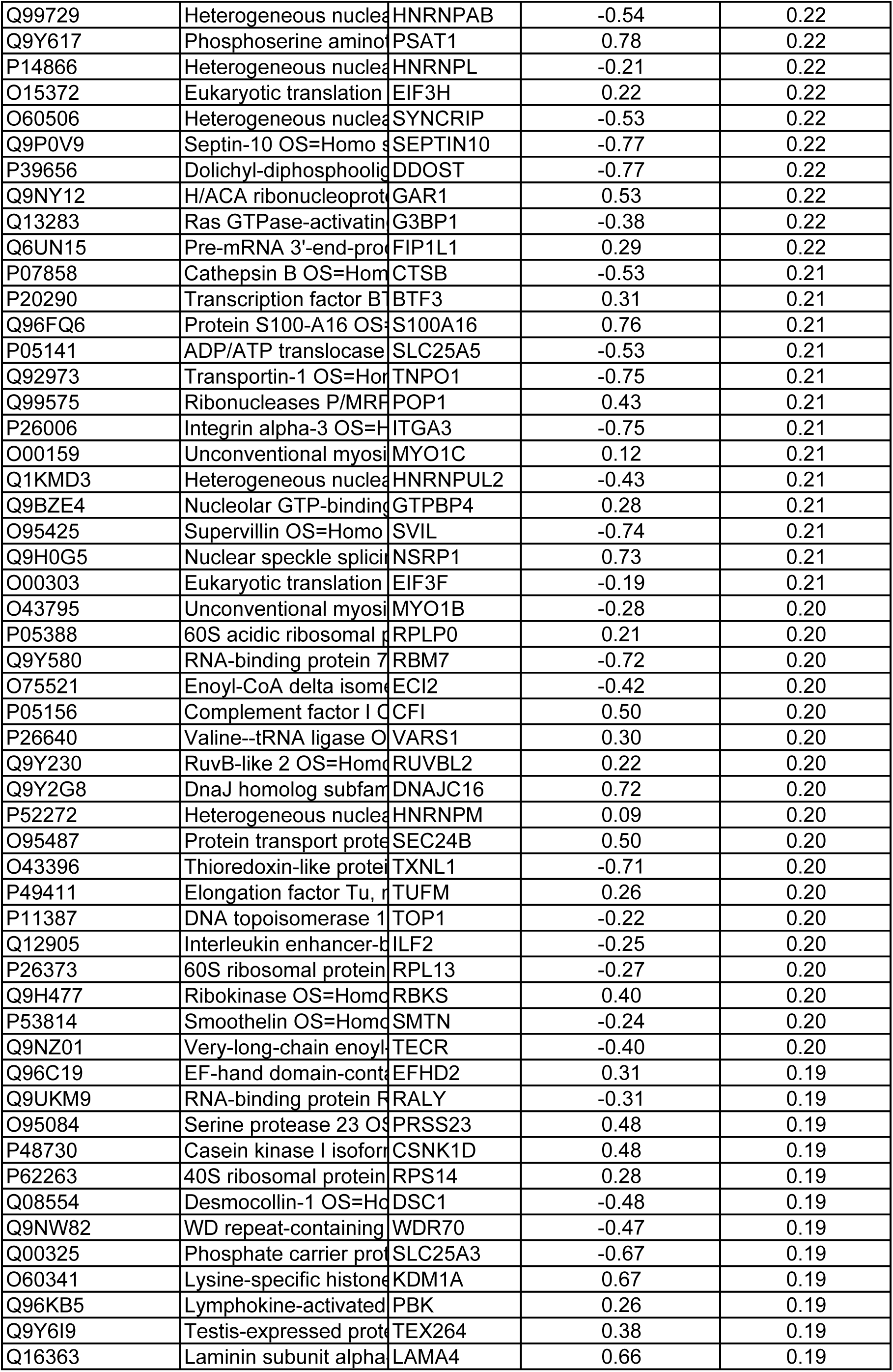
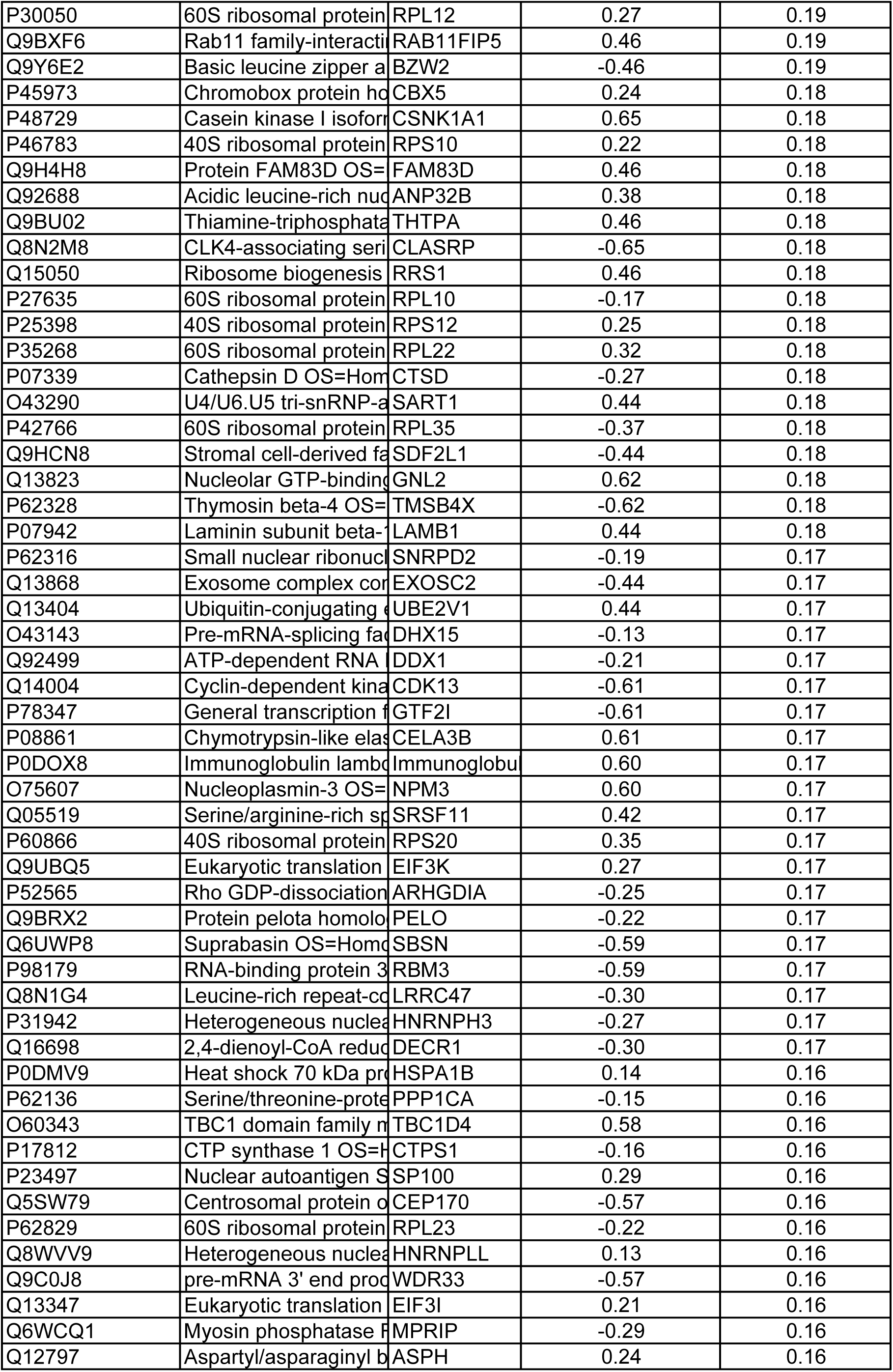
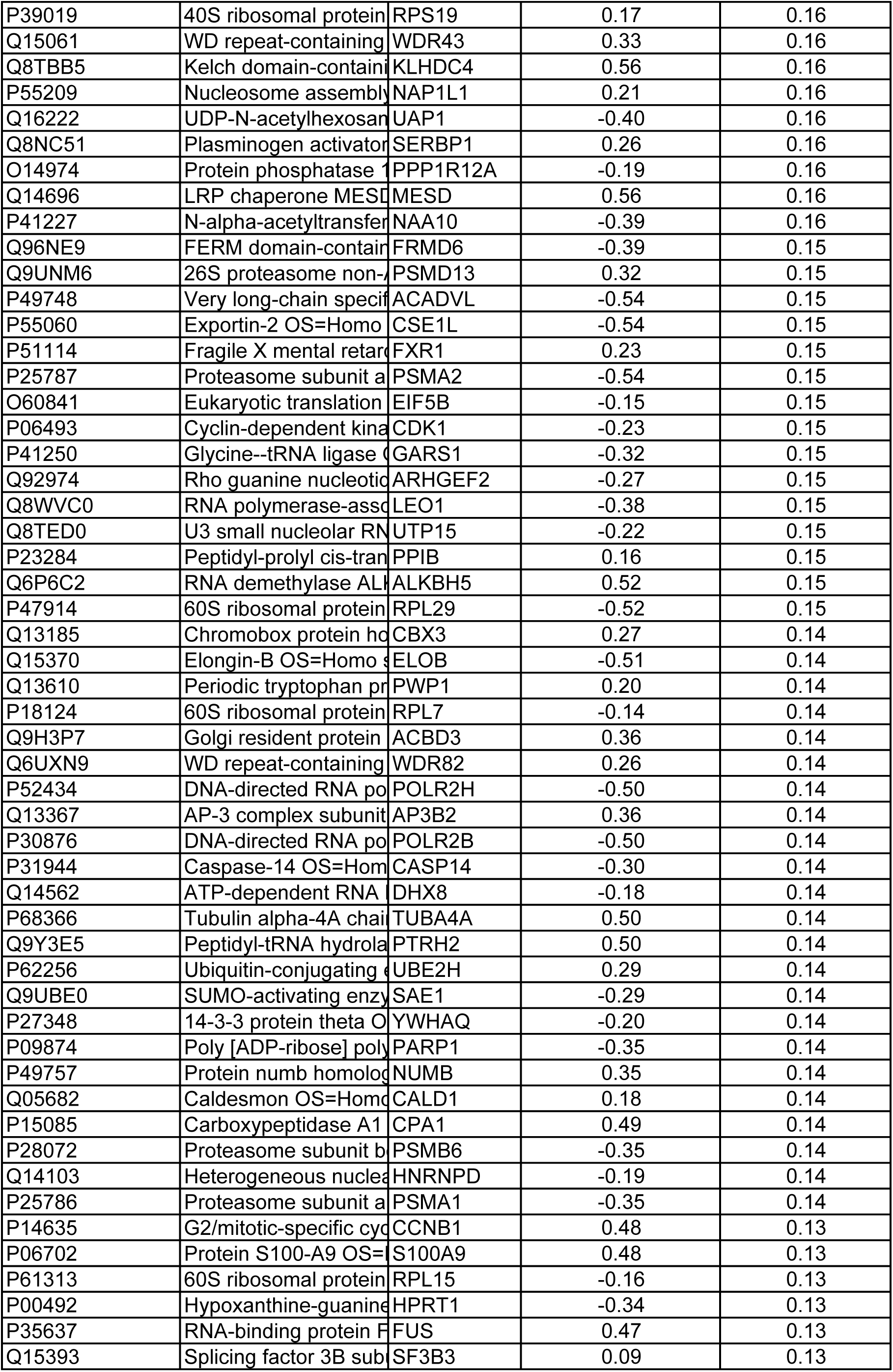
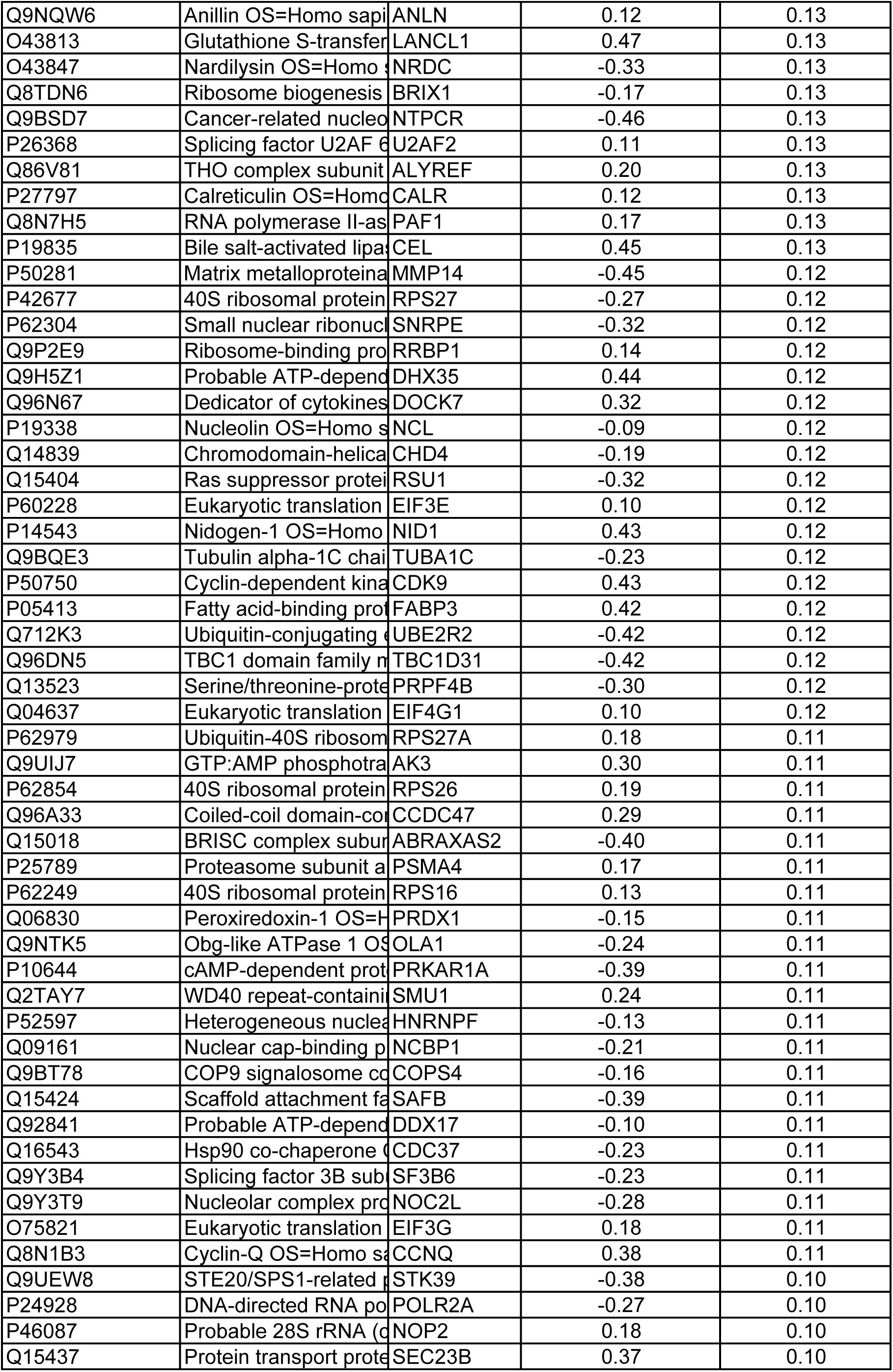
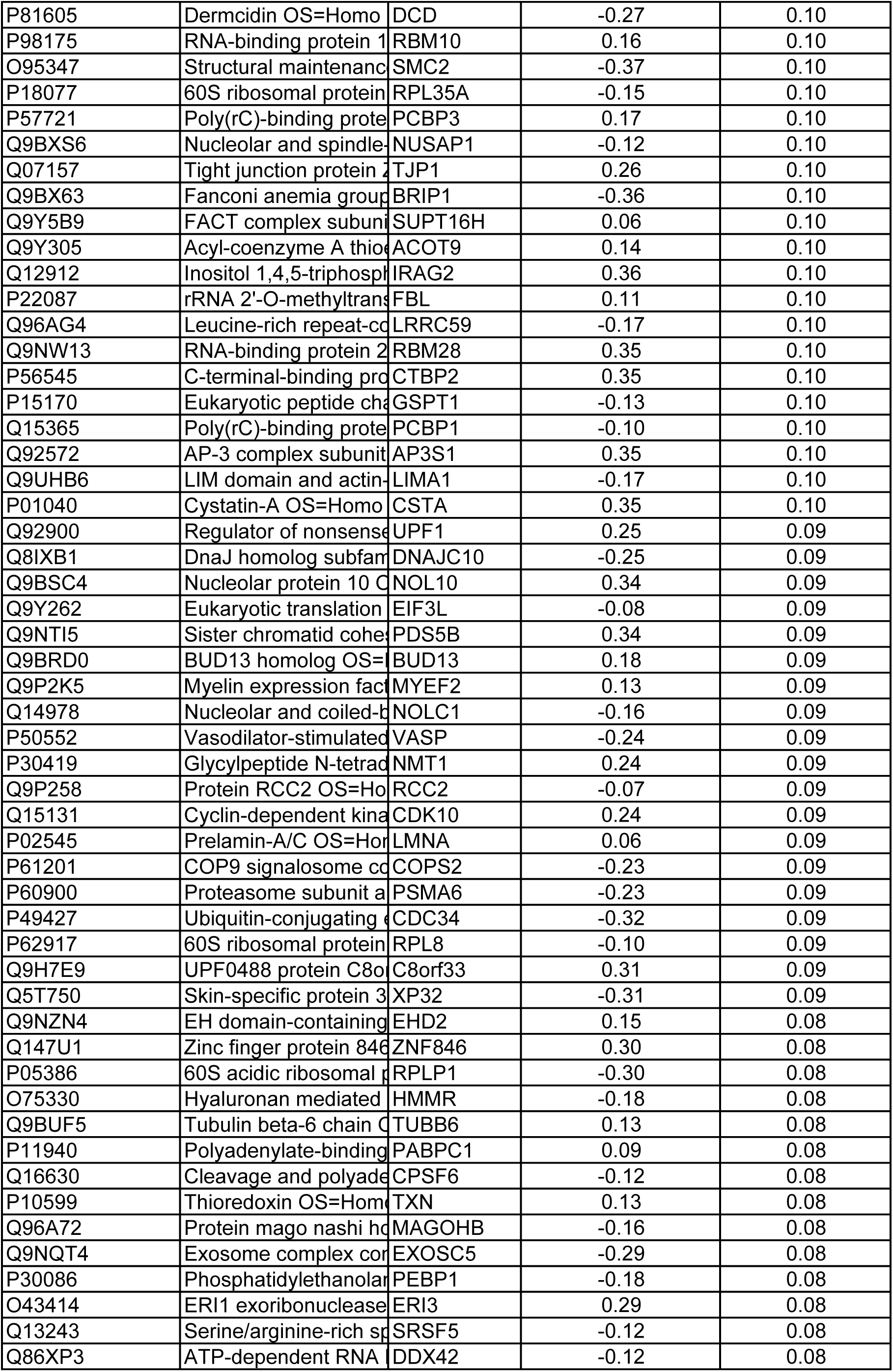
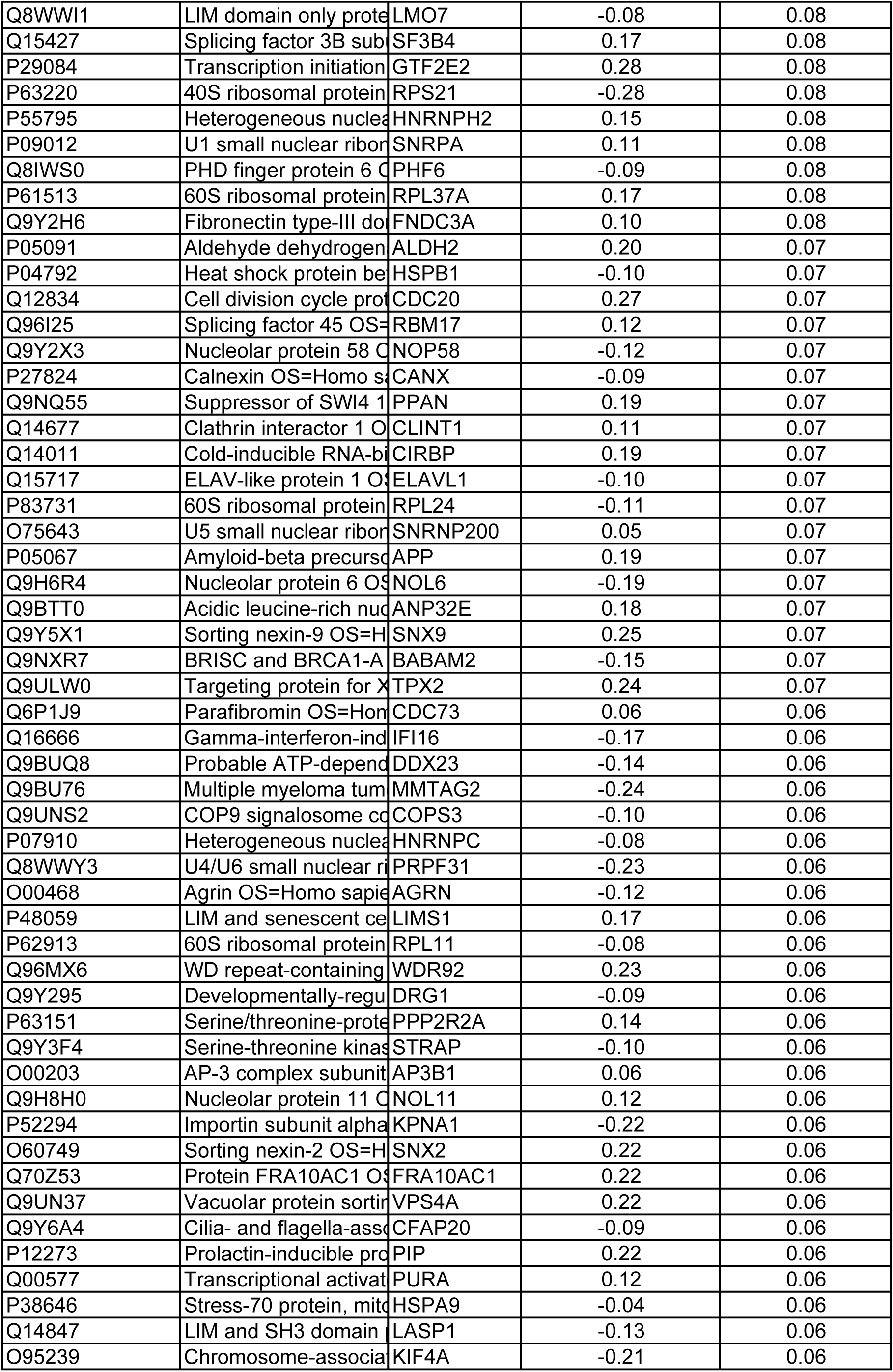
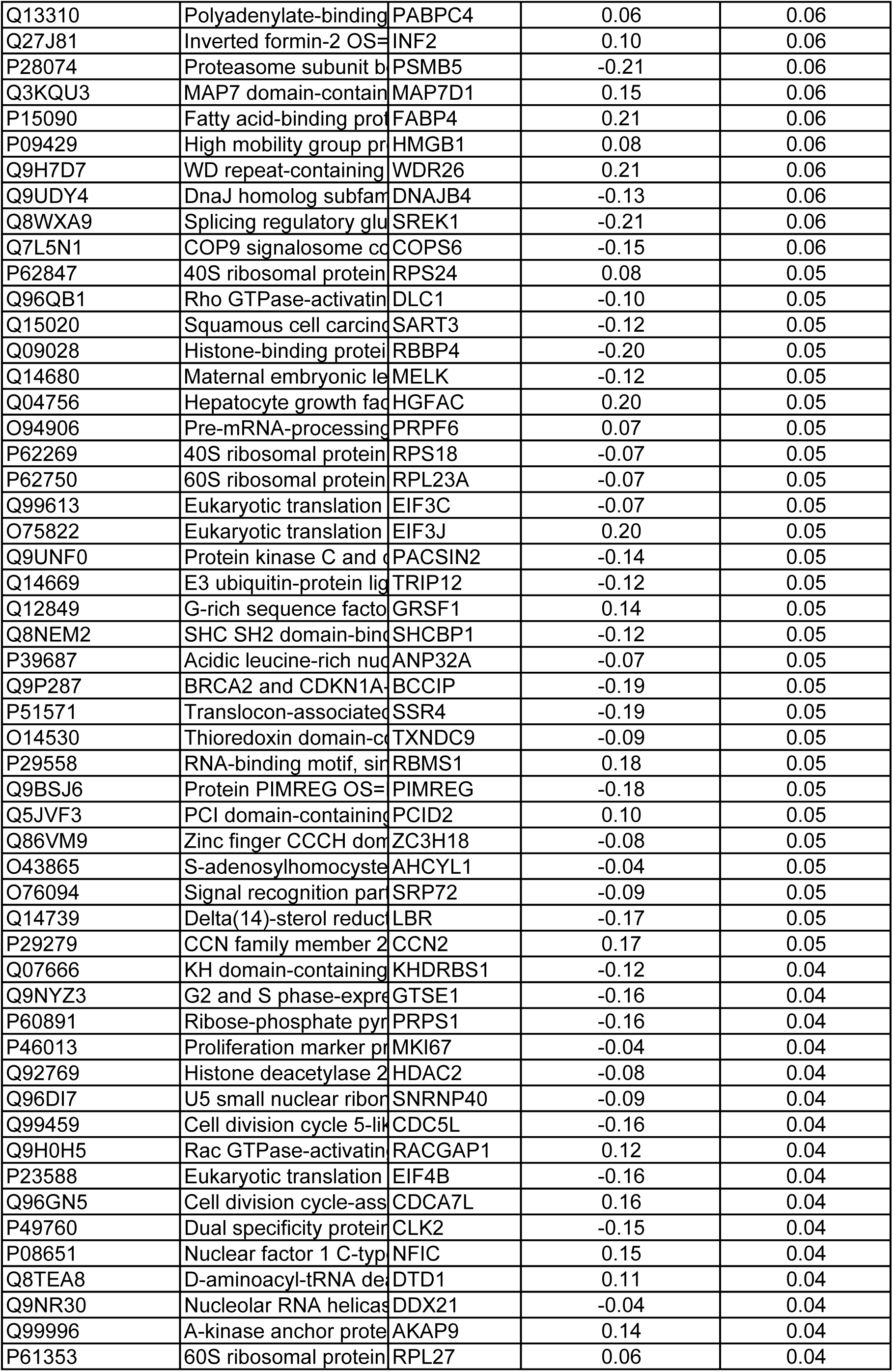
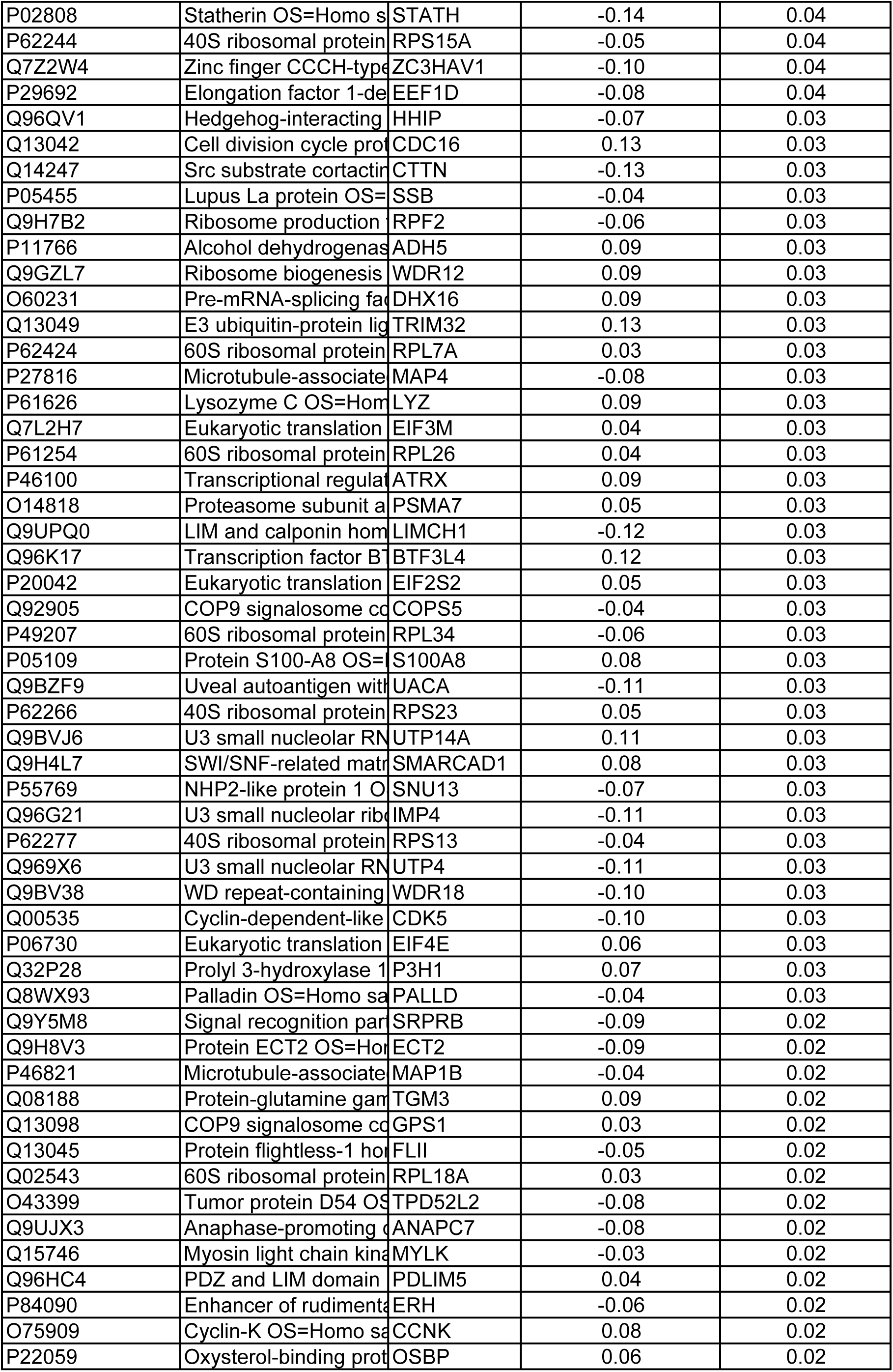
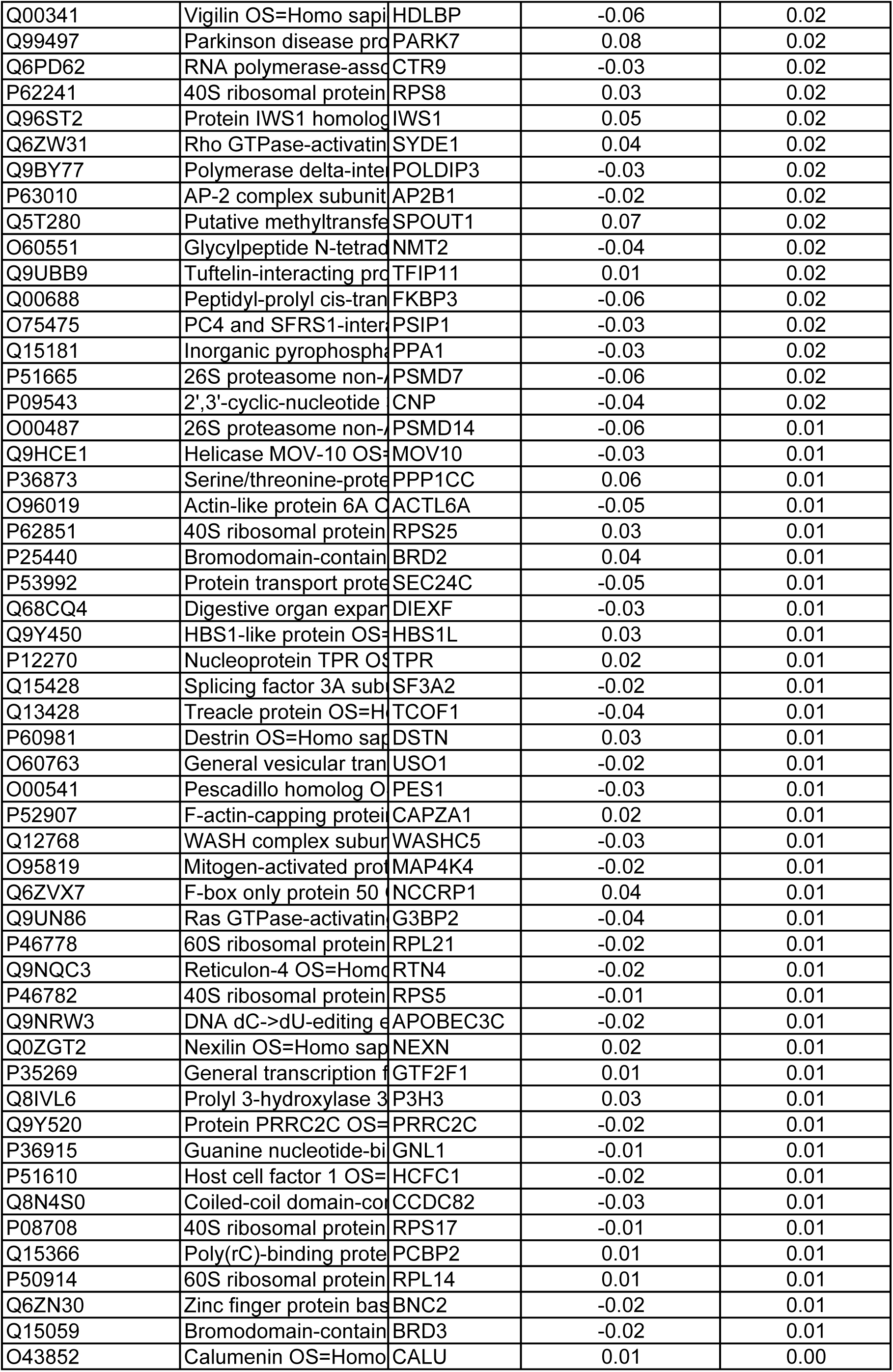
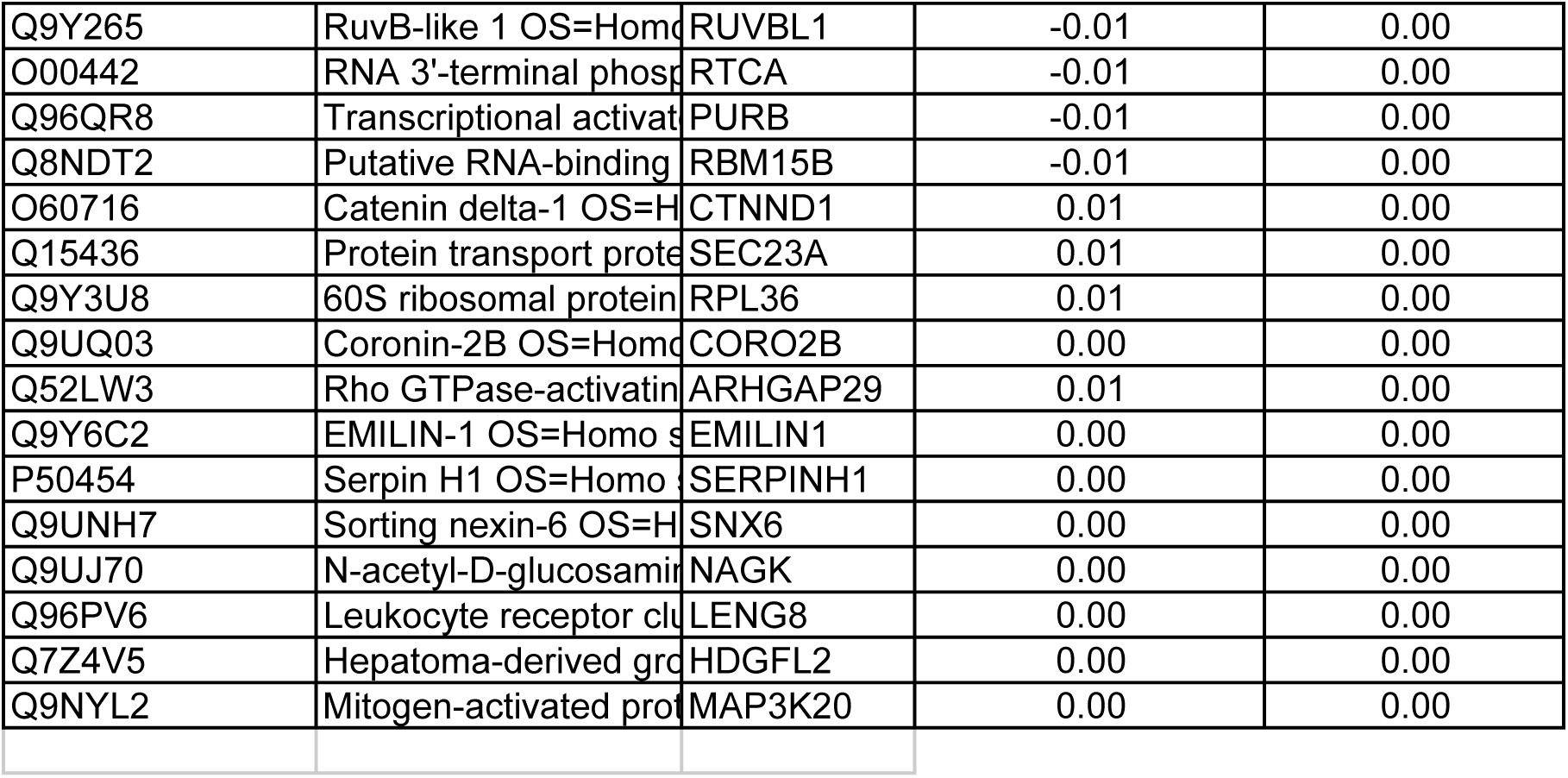
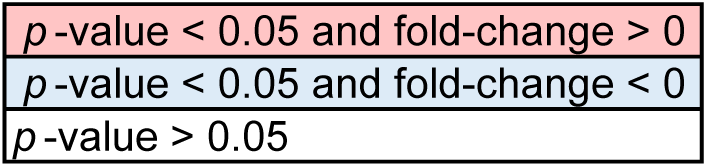
The table lists the 53BP1 mitotic interactors enriched in the V5 pull-downs from RPE1 53BP1-V5 cells as compared to RPE1 WT cells.

Next, we investigated whether CENP-F is necessary for 53BP1 recruitment to the KT. siRNA-mediated depletion of CENP-F in HeLa S3 cells disrupted 53BP1 localization at mitotic KTs (Fig. 1E-F), without affecting 53BP1 protein levels (Fig. EV1D). In order to tackle whether the abovementioned findings can be recapitulated in a non-transformed, p53-proficient cell line we performed CENP-F depletion in RPE1 cells. Strikingly, although 53BP1 localization to KTs appeared dependent on CENP-F also in this setting, releasing 53BP1 in the cytosol did not appear to alter the MSP threshold in RPE1 cells (Fig. 1G-H).

Taken together, our data support the notion that CENP-F binds to and recruits 53BP1 to the KT. However, the functional relevance of this localization remains to be defined.

### A CENP-F point mutant separates CENP-F functions and displaces 53BP1 from the KT

Although *CENPF* knock-out HeLa cells have been reported to display normal growth (McKinley & Cheeseman, 2017), CENP-F is critical for assisting KT-microtubule attachment and protecting corona cargoes from dynein dependent “stripping” (Bomont *et al*, 2005; Auckland *et al*, 2020). Consistent with this notion, the median Chronos gene score (Dempster *et al*, 2021) for *CENPF* knock-out in 1078 different cell lines analysed in Depmap was -0.23, and the gene appeared essential in 113 of them (Dempster *et al*, 2019), demonstrating that complete knock-out of *CENPF* confers a loss of fitness (Fig. 2A). Therefore, we sought a targeted strategy to inhibit CENP-F binding to 53BP1. Because of the high number of CENP-F clones retrieved by our yeast two-hybrid screen, we were able to restrict the CENP-F minimal interacting domain with 53BP1 to a region of 25 residues, aa 564-588 (Fig. 2B). Molecular modelling revealed that this domain resides in a region of the protein predicted to be part of a coiled-coil motif (Ciossani *et al*, 2018). Thus, substituting the glutamic acid (E) at position 564 with proline (P, which destabilizes the α-helical fold, Fig. 2C), could provide a strategy to selectively hinder protein-protein interactions relying on the proper folding of this coiled-coil region.

**Figure 2.**
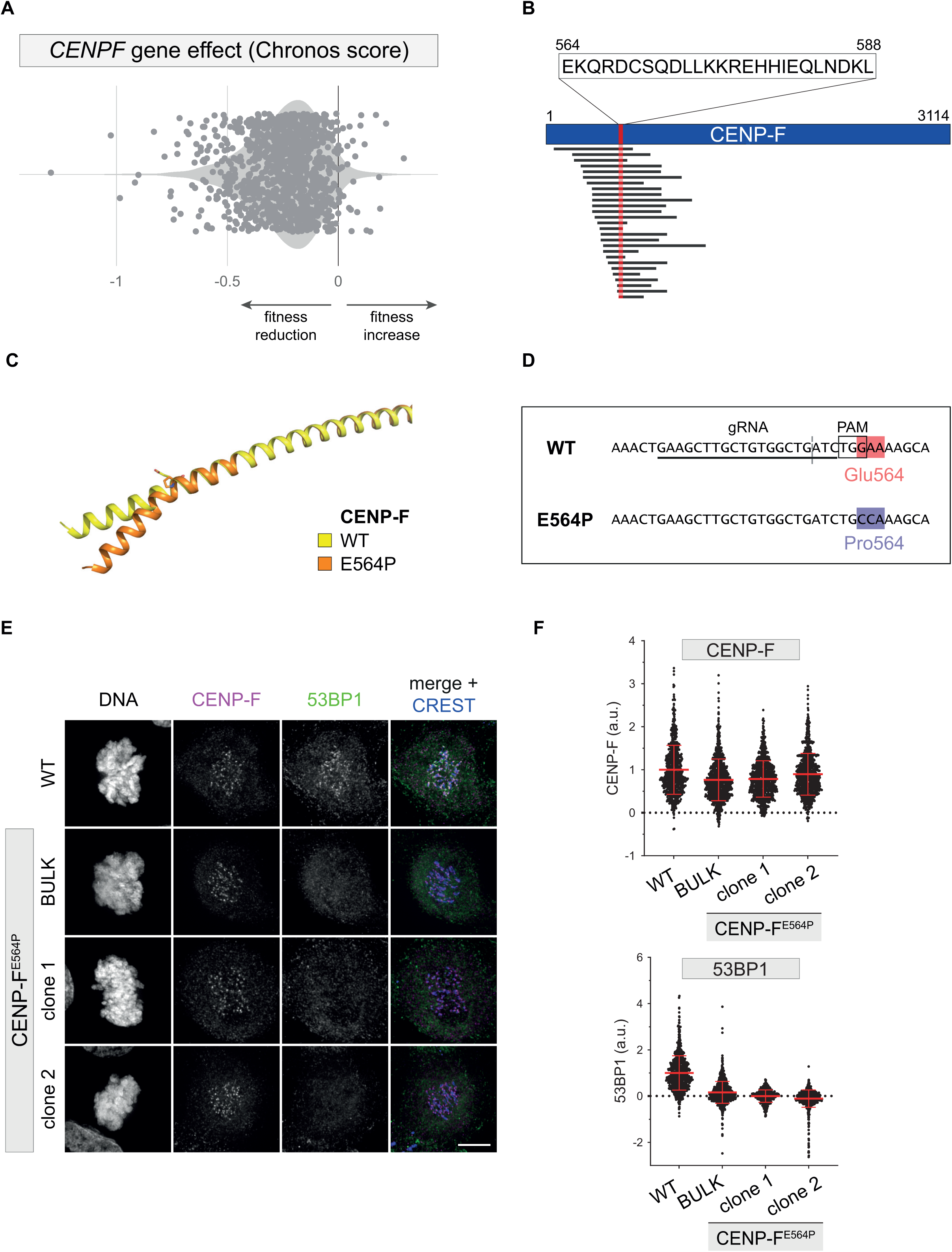
CENP-F^E564P^ displaces 53BP1 from the kinetochore. A) Chronos gene dependency score of *CENPF* across 1078 cell lines from DepMap database (CRISPR DepMap 22Q4). X-axis: Chronos score. B) Schematic of the CENP-F prey fragments retrieved by the yeast two-hybrid screen and their relative position to the CENP-F sequence. The overlap between all the different clones identifies a region of 25 amino acids as putative binding domain between the KT-binding domain of 53BP1 (bait) and CENP-F. C) AlphaFold modelling of the putative 53BP1 binding domain of CENP-F. Introduction of a proline in position 564 is predicted to introduce a kink in this coiled coil region. D) Schematic depicting the knock-in strategy used to introduce E564P mutation in *CENPF*. The gRNA recognition site, PAM sequence and cut site (dashed vertical line) are presented. E) Fluorescent micrographs of RPE1 cells of the indicated genotype, co-stained with the indicated antibodies. Scale bar: 5 µm. F) Dot plots showing the intensity of the indicated proteins at individual KTs. Mean values (red lines) ± SD are reported. Data obtained from images as in D). N ≥ 462 KTs were assessed from at least 20 cells for each condition; a.u. = arbitrary units.

To this end, we used CRISPR/Cas9 to introduce the E564P substitution into the endogenous *CENPF* locus of RPE1 cells (Fig. 2D) (Ghetti *et al*, 2021). One caveat of this experiment is that, if effective in delocalizing 53BP1 from KTs, the CENP-F^E564P^ mutation might lead to hyperactivation of the MSP, promoting cell cycle arrest. This effect could, in turn, lead to difficulty in expanding the population of edited cells. Strikingly, however, sequencing of the electroporated polyclonal population (hereafter referred to as bulk) confirmed the introduction of the desired mutation with a high editing rate at 5 days post-electroporation (that is, > 80%, Fig. EV2A). The bulk population re-sequenced on day 15 revealed that penetrance of the desired edit did not decrease over time (Fig. EV2A). This shows that under our experimental conditions, the CENP-F^E564P^ mutation is compatible with cell proliferation in RPE1 cells. Moreover, homozygous monoclonal CENP-F^E564P^ RPE1 derivatives were obtained from the polyclonal population (Fig. EV2A).

CENP-F^E564P^ mutant cells proved to be devoid of 53BP1 recruitment to mitotic KTs, phenocopying CENP-F depletion (Fig. 2E-F). Importantly, however, this point mutation did not abolish CENP-F recruitment to the corona (Fig. 2E-F) or its ability to recruit the downstream effector protein NudE (Fig. EV2B-C). Thus, we precisely mapped the CENP-F minimal domain required for 53BP1 recruitment to the KT and demonstrated that the introduction of a single amino acid substitution in the CENP-F sequence allowed the generation of a separation-of-function mutant, which interferes with 53BP1 binding without impairing its ability to localize at the KT and recruit other downstream components such as NudE.

### 53BP1 KT localization is dispensable for MSP functionality

The generation of CENP-F^E564P^ mutant cell lines provided a unique tool to precisely dissect the functional relevance of 53BP1 KT localization. We reasoned that 53BP1 recruitment at the fibrous corona could reflect a priming state for the MSP, similar to what happens for SAC proteins recruited to the same structure. However, we obtained strong evidence against this hypothesis. Activating the MSP with centrinone, a specific PLK4 kinase inhibitor causing centriole depletion over time, led to a measurable drop in the clonogenic potential of wild type (WT) RPE1 cells (Fig. 3A). Expectedly, MSP-deficiency (achieved by *TP53BP1* KO) clearly boosted the clonogenic potential in the presence of centrinone. Strikingly, however, this behaviour was not matched in CENP-F^E564P^, neither when utilizing the bulk population nor the homozygote clonal derivatives (Fig. 3 A-B). The same picture emerged when evaluating the proliferation rate displayed by the various genotypes in the presence of centrinone: competition-based growth assays highlighted that *TP53BP1* KO was the only genotype capable of weakening the cell cycle arrest promoted by centrinone, while CENP-F^E564P^ mutant cells displayed an arrest that was at least as proficient as the one observed in WT cells (Fig. 3C). Moreover, the functionality of the MSP in CENP-F^E564P^ knock-in cell lines could also be verified using a different trigger of the pathway, namely SAC activation, transiently exposing mitotic cells to nocodazole, promoting an extension of mitotic duration. Also in such experimental conditions, *TP53BP1* KO was the only genotype retaining high clonogenic potential, wherease extension of mitotic duration in CENP-F^E564P^ knock-in cell lines triggered a reduction of the clonogenic potential that was at least as pronounced as in WT cells (Fig. 3 D-E). Taken together, our data demonstrate that 53BP1 recruitment at the KT is not necessary for MSP function.

**Figure 3.**
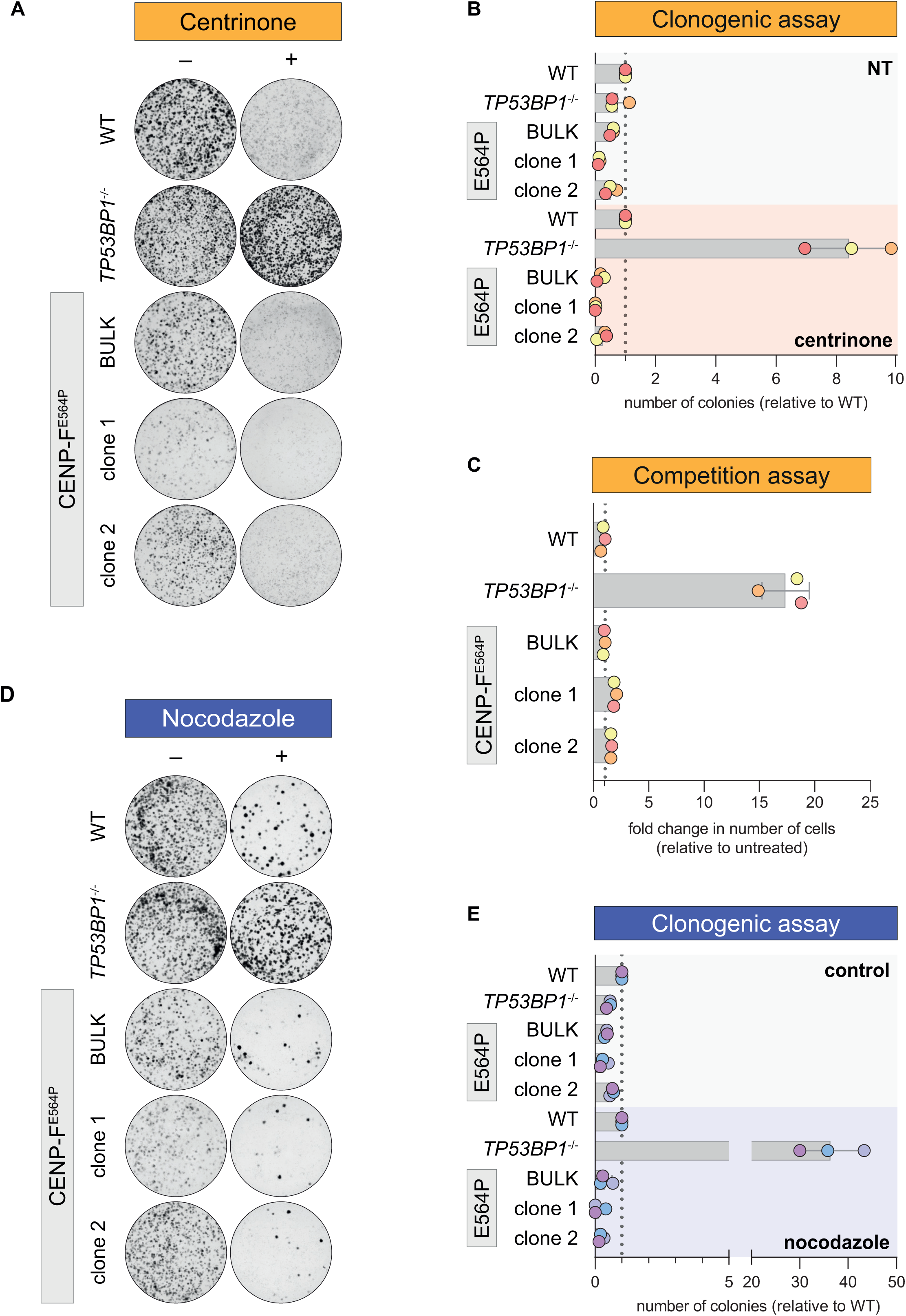
53BP1 kinetochore localization is dispensable for MSP functionality. A) Cells of the indicated genotype were treated with centrinone for 12 days or left untreated (for 8 days) and stained with crystal violet. Images are representative of N = 3 biologically independent experiments. B) Quantification of 3 independent experiments as in A). Data are presented as number of colonies relative to WT cells. The bars indicate the mean ± SD (N = 3 independent experiments). C) Cells of the indicated genotype were mixed 1:1 with EGFP-expressing WT cells and cultured either in the presence or absence of centrinone. After 8 days, the relative growth of each genotype was assessed by flow cytometry. The bars indicate the mean ± SD (N = 3 independent experiments). D) Cells of the indicated genotype were treated with thymidine for 24 h and released in medium containing 3.3 µM nocodazole for 16 h. Mitotic cells were selectively retrieved by shake-off, seeded for the clonogenic assay and stained with crystal violet after 12 days. Control cells were synchronized by a thymidine block, released in fresh medium, detached by trypsinization after 16 h, seeded for the clonogenic assay, and stained after 8 days. Images are representative of N = 3 biologically independent experiments. E) Quantification of 3 independent experiments as in D). Data are presented as number of colonies relative to WT cells. The bars indicate the mean ± SD (N = 3 independent experiments).

### PLK1 promotes 53BP1 loss of KT affinity and MSP activation

While our CENP-F mutant cell lines afforded a way to displace 53BP1 from the KT, it was unclear whether it is possible to perturb the system in the opposite manner, namely forcing 53BP1 to remain at the KT. To gain better insight into this aspect, we characterized 53BP1 behaviour during mitosis in greater detail.

In agreement with previous studies (Jullien *et al*, 2002), 53BP1 displayed maximal association with mitotic KTs during prophase and then gradually dissociated in prometaphase and the subsequent phases (Fig. EV1C). This localization pattern was similar to that reported for other corona-localizing proteins. In fact, in the context of normal mitosis, the outermost fibrous corona layers are disassembled via dynein-motor protein activity upon microtubule-KT attachment, a process called stripping (Auckland *et al*, 2020). In a stripping hyperactivation assay (Howell *et al*, 2001), 53BP1 was removed from the KT and accumulated at the spindle poles (Fig. EV3A), suggesting that, likewise CENP-F (Auckland *et al*, 2020), 53BP1 is also a dynein cargo during mitosis. In this perspective, it is not surprising that the amount of 53BP1 detected at the KT decreased over time in the presence of a monoastral spindle caused by Eg5 inhibition (Fong *et al*, 2016). Strikingly, however, in the presence of the microtubule-poison nocodazole, the 53BP1 signal was still gradually lost over time, completely disappearing from KTs at around 6 h after mitotic entry (Fig. EV3B). This suggests that in addition to stripping, 53BP1 is subjected to a time-dependent loss of affinity for the KT.

Phosphorylation events play crucial roles in mitotic KT assembly, dynamics, and remodelling (Musacchio & Desai, 2017; Saurin, 2018). Moreover, 53BP1 is known to be hyperphosphorylated during mitosis (Jullien *et al*, 2002; Lee *et al*, 2014; Giunta *et al*, 2010). Therefore, we assessed whether the major known kinases targeting 53BP1 (i.e., ATM, ATR, PLK1, and Aurora B), along with master regulator kinases acting at the KT (MPS1 and BUB1), contribute to modulating the affinity of 53BP1 for the KT. Our mini-screen showed that inhibition of none of the tested kinases abolished 53BP1 loading to KTs, although ATR inhibition caused a measurable loading reduction (Fig. 4A). In a complementary setting, we also evaluated the contribution of the aforementioned kinases to 53BP1’s loss of affinity for the KT. This analysis revealed that PLK1 has a clear contribution, as its inhibition alters 53BP1 localization dynamics, abolishing the time-dependent loss of affinity for the KT (Fig. 4B). To exclude any potential off-target effect of the PLK1 inhibitor used in our mini-screen, we took advantage of RPE1 cells in which endogenous PLK1 has been deleted and rely on the constitutive expression of a genetically modified allele (PLK1^AS^). This allele can be chemically inactivated by a bulky purine analogue which occupies its ATP-binding pocket but is inactive toward cells lacking the PLK1^AS^ allele (Burkard *et al*, 2007). Also in this setting, inhibition of PLK1 in the context of MSP activating conditions was able to retain 53BP1 KT localization in WT cells but it did not restore the impaired localization of the protein in our CENP-F^E564P^ mutant (Fig. 4C and EV3C).

**Figure 4.**
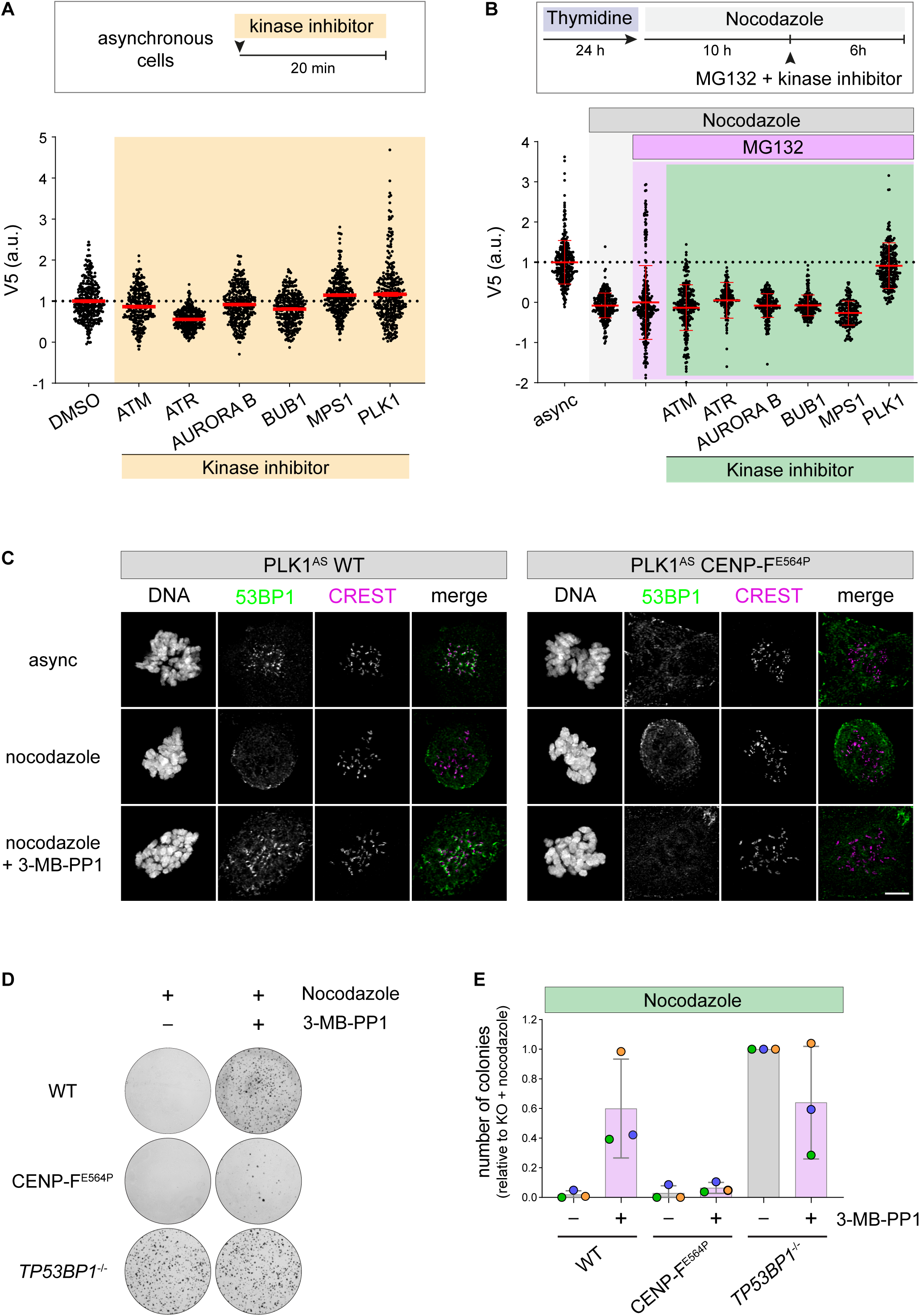
PLK1 activity is required for the MSP. A) Top: Asynchronous RPE1 53BP1-V5 cells were treated for 20 min with the specific kinase inhibitor. Bottom: dot plots showing V5 fluorescence intensity at individual KTs. Mean values (red lines) ± SD are reported. N ≥ 259 KTs were assessed from at least 20 cells for each condition; a.u. = arbitrary units. B) Top: RPE1 53BP1-V5 cells were treated with thymidine for 24 h and released in medium containing 3.3 µM nocodazole. After 10 h, MG-132 and the specific kinase inhibitor were added. Bottom: dot plots showing V5 fluorescence intensity at individual KTs. Mean values (red lines) ± SD are reported. N ≥ 165 KTs were assessed from at least 20 cells for each condition; async = asynchronous cells; a.u. = arbitrary units. C) RPE1 PLK1^AS^ cells of the indicated genotype were treated with thymidine for 24 h and released into medium containing 3.3 µM nocodazole for 16 h, in the presence or absence of PLK1 inhibition (3-MB-PP1). Representative fluorescence micrographs are shown. Scale bar: 5 µm. D) RPE1 PLK1 ^AS^ cells of the indicated genotype were treated as in C). Mitotic cells were selectively retrieved by shake-off, washed-out from the drugs, seeded for the clonogenic assay, and stained with crystal violet after 16 days. Images are representative of N = 3 biologically independent experiments. E) Quantification of 3 independent experiments as in D). Data are presented as number of colonies relative to the *TP53BP1*^-/-^ + nocodazole condition. The bars indicate the mean ± SD (N = 3 independent experiments).

The PLK1 contribution in 53BP1 KT loss of affinity prompted us to evaluate whether the maintenance of 53BP1 at the KT via PLK1 inhibition has an impact on MSP functionality. To this end, we synchronized PLK1^AS^ WT and CENP-F^E564P^ cells in mitosis and artificially prolonged prometaphase in the presence or absence of the ATP analogue. We then released the cells and evaluated their long-term proliferative capabilities by assessing their clonogenic potential. Strikingly, PLK1 inhibition restored cell cycle re-entry in WT cells, demonstrating that PLK1 activity during extended prometaphase is crucial for MSP function (Fig. 4D-E). Given that PLK1 inhibition during the transient exposure of the cells to nocodazole appears sufficient to retain 53BP1 at the KT and to promote MSP override, we exploited the CENP-F^E564P^ mutant to assess whether these two phenomena were causally linked. Strikingly, when PLK1 inhibition was performed in cells in which 53BP1 loading to KT was prevented due to the CENP-F^E564P^ mutation, the MSP override observed in WT cells was also abrogated (Fig. 4D-E). Taken together, our data demonstrate that PLK1 activity supports MSP functionality by promoting a time-dependent loss of affinity of 53BP1 from the KT.

In the present study, we addressed the functional relevance of 53BP1 KT localization for the MSP. The fact that 53BP1 dynamically associates with KTs has been known for over two decades (Jullien *et al*, 2002). On the other hand, the contribution of 53BP1 to the MSP was solidly demonstrated in 2016 (Fong *et al*, 2016; Lambrus *et al*, 2016; Meitinger *et al*, 2016). However, how a mitotic delay can be translated into stimulation of USP28 enzymatic activity towards the p53 protein remained enigmatic. Here, we show that 53BP1 physiologically associates with KTs thanks to a direct interaction with the fibrous corona protein CENP-F. Surprisingly, this association appeared dispensable for MSP proficiency. Moreover, we demonstrated that time-dependent loss of affinity of 53BP1 for the KT is a phenomenon that can be perturbed by PLK1 enzymatic inhibition. PLK1 is known to associate with outer KTs (Singh *et al*, 2021) and to phosphorylate both CENP-F and 53BP1 (van Vugt *et al*, 2010; Santamaria *et al*, 2010). Thus, it appears likely that local direct phosphorylation of either protein concurs to 53BP1 KT loss of affinity (Fig. 5).

**Figure 5.**
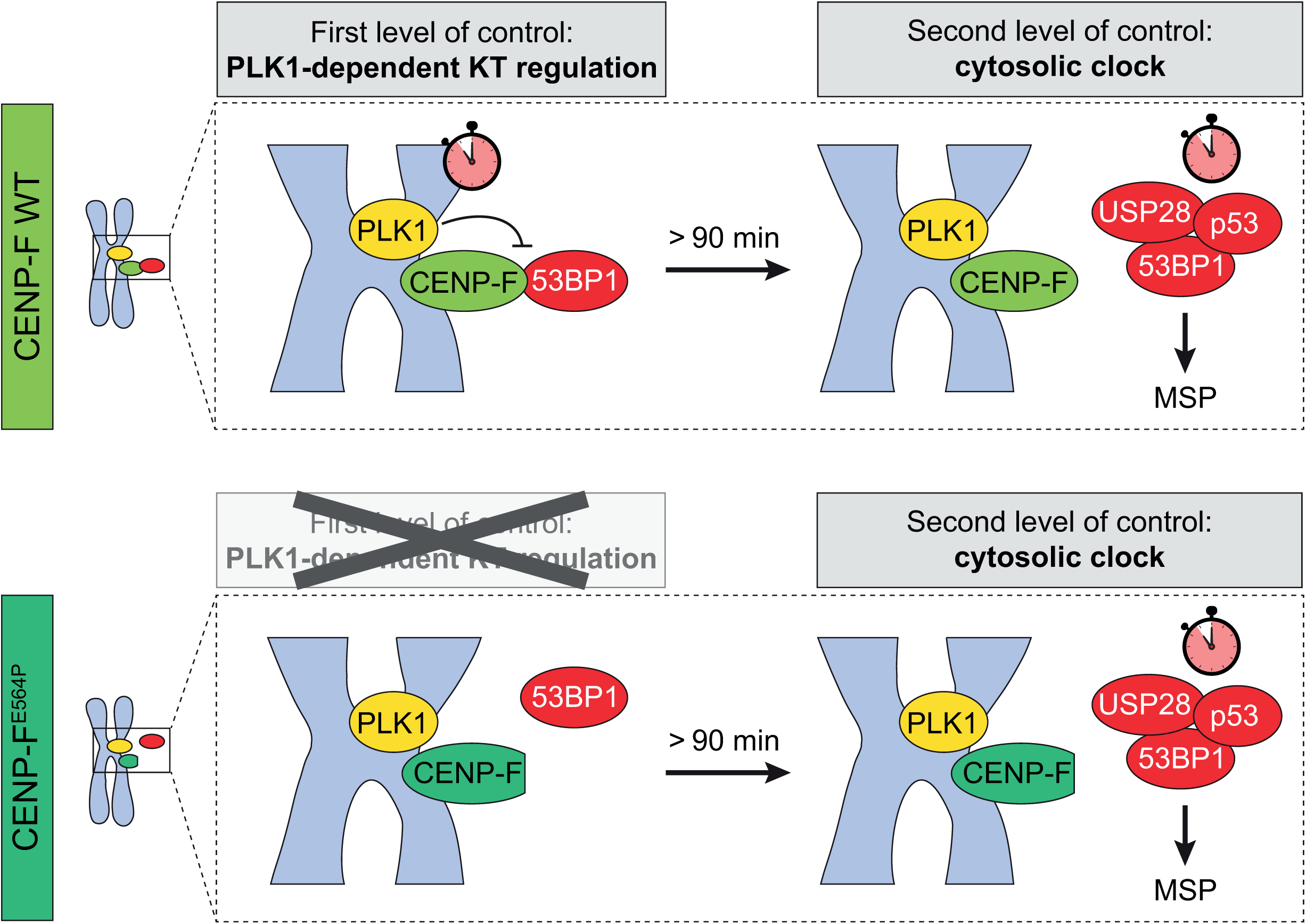
The MSP relies on a belt and braces regulation. In a normal mitosis, 53BP1 is recruited to the KT via a direct interaction with CENP-F. In the context of a prolonged mitosis, PLK1 activity promotes the loss of affinity of 53BP1 for the KT (first priming step). Cytosolic 53BP1 needs an additional yet unidentified trigger to induce the USP28-dependent deubiquitination of p53 which leads to cell cycle arrest (second priming step). Interfering with 53BP1 KT localization bypasses the PLK1-dependent regulatory layer maintaing MSP proficiency.

Importantly, PLK1 inhibition also leads to MSP abrogation. Crucially, our CENP-F^E564P^ mutation highlights that when 53BP1 is no longer capable of KT association, PLK1 appears no longer necessary for MSP proficiency (Fig. 5). Thus, while we demonstrated that PLK1 contribution to the MSP functionality relies on 53BP1 KT displacement, it is also clear that a yet to be identified cytosolic mechanism allows the measurement of mitotic timing. In this perspective, PLK1-dependent KT release of 53BP1 might not represent the central mitotic timer, yet we speculate that this regulatory layer might concur to the generation of a stringent and reproducible timer threshold. Finally, we hypothesize that other cytosolic mechanisms, e.g. the regulation of 53BP1 propensity to form higher order oligomers and to undergo liquid-liquid phase separation (Kilic *et al*, 2019), might concur to determine the ability to measure mitotic timing and to transduce it to the p53 tumor suppressor pathway.

## Materials and Methods

### Cell culture

hTERT-RPE1 cells (a gift of Stephan Geley, Medical University of Innsbruck), hTERT-RPE1 PLK1^AS^ cells (a gift of Prasad Jallepalli, Memorial Sloan Kettering Cancer Center) and hTERT RPE1 p21-EGFP cells (a gift of Chris Bakal, Institute of Cancer Research) were cultured in DMEM/F12 1:1 (Gibco). HEK 293T (a gift from Ulrich Maurer, University of Freiburg) and HeLa S3 (a gift from Erich Nigg) cells were maintained in DMEM (Gibco). All media were supplemented with 10% foetal bovine serum (Gibco), 2 mM L-glutamine (Corning), and penicillin-streptomycin solution (Corning). Cells were grown at 37°C with 5% CO_2_ and routinely tested for mycoplasma contamination.

### Drug treatments, irradiation and synchronization procedure

The following compounds were used: 1 µM NU-7441 (Selleck Chemicals), 125 nM centrinone (MedChem Express), 2 mM thymidine (Abcam), 3.3 µM nocodazole (BioTrend), 2 µM dimethylenastron (Selleck Chemicals), 100 nM BI-2536 (MedChem Express), 500 nM reversine (Enzo Life Sciences), 10 µM KU-60019 (MedChem Express), 10 µM VE-822 (MedChem Express), 2 µM ZM-447439 (Selleck Chemicals), 10 µM BAY-524 (MedChem Express), 10 µM MG-132 (Selleck Chemicals), 10 µM 3-MB-PP1 (Cayman Chemical). Thymidine was dissolved in water whereas all other drugs were dissolved in DMSO; to all untreated controls solvent only was administered. Cells were irradiated with 2 Gy (delivering at a dose rate of 1.6 Gy/min) of ionizing radiation using the Xstrahl RS225 X-ray research Irradiator (West Midlands, UK) and allowed to recover for 30 min prior to fixation for immunofluorescence. Synchronization of RPE1 was performed by arresting cells with thymidine for 24 h, followed by release in fresh medium containing nocodazole for either 10 h or 16 h. Mitotic cells were then harvested by selective shake-off and either directly lysed or washed four times and released into fresh medium. For the ATP depletion assay (Howell *et al*, 2001), cells were rinsed in saline (140 mM NaCl, 5 mM KCl, 0.6 mM MgSO_4_, 0.1 mM CaCl_2_, 1 mM Na_2_HPO_4_, 1 mM KH_2_PO_4_, pH 7.3) and placed in either saline G (saline + 4.5 g/l D-glucose) or Az/DOG (saline + 5 mM sodium azide + 1 mM 2-deoxy-D-glucose) for 15 min at 37°C.

### siRNA-mediated gene knock-down

HeLa cells were transfected with 40 nM CENP-F siRNA (5’-CAAAGACCGGUGUUACCAAG-3’) or luciferase (GL2) siRNA (5’-CGUACGCGGAAUACUUCGA-3’) whereas RPE1 cells where transfected with siRNA SMART pool targeting CENP-F (Dharmacon, L-003253-00-0005) or a non-targeting control (Dharmacon, D-001810-01-05) using Lipofectamine RNAiMAX Transfection Reagent (Life Technologies), according to the manufacturer’s procedures. 48 h post-transfection, cells were either harvested for Western blotting or fixed for immunofluorescence analysis.

### AlphaFold molecular modelling

CENP-F models for WT and the E564P mutant were obtained through ColabFold v1.5.2 (Mirdita *et al*, 2022). Sequence boundaries (aa 546-780) were chosen to include all residues predicted in the AlphaFoldDB (Jumper *et al*, 2021) in the single α-helix comprising Glu564.

### Engineering of *TP53BP1* and *CENPF* endogenous loci

2×10^5^ RPE1 cells were electroporated using Lonza Nucleofector 4-D according to manufacturer’s instruction and as previously described (Ghetti *et al*, 2021). Briefly, 100 μM of crRNA (IDT) and 100 μM of tracRNA (IDT) were annealed to form gRNAs. Ribonucleoparticles (RNPs) were complexed by mixing 120 pmol of recombinant Cas9 and 150 pmol of gRNAs. Electroporation mix was prepared resuspending RPE1 cells in P3 Primary Cell Full Electroporation Buffer (Lonza), and adding previously prepared RNPs, 4 μM Alt-R Cas9 Electroporation enhancer (IDT) and 4 μM single-strand DNA homology template (Ultramer DNA Oligonucleotide, Lonza). After electroporation, cells were treated with 1 μM NU-7441 (Selleck Chemicals) for 48 h and single-cell clones were obtained by dilution cloning. Genomic DNA was extracted using NucleoSpin Tissue columns (Macherey-Nagel). Sanger sequencing of PCR products spanning the insertion sites confirmed the correct in-frame insertion of the V5 tag (*TP53BP1* locus) or E564P mutation (*CENPF* locus) in homozygosis. For crRNA sequences, DNA homology donors and PCR primers see Table 2.

**Table 2.**
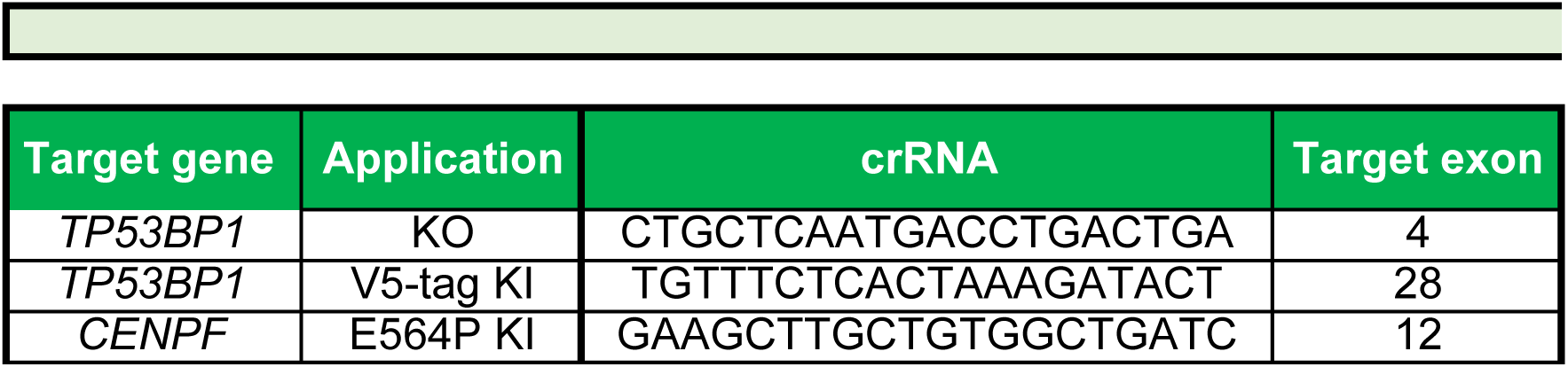
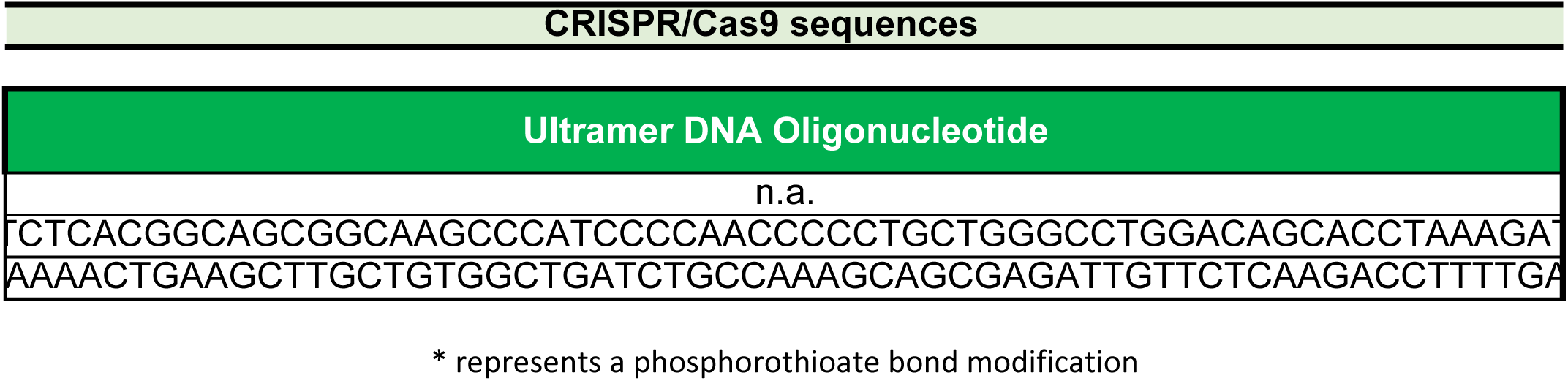
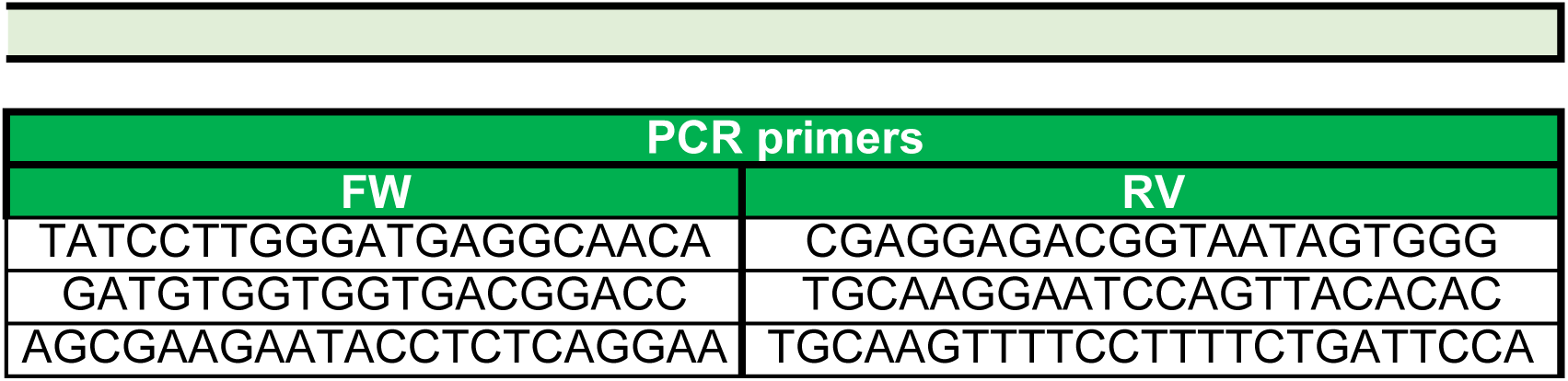
The table lists the gRNA, Ultramer DNA oligonucleotide homology template, and PCR primer sequences used in this study.

### Generation of *TP53BP1* KO cell lines

RPE1 *TP53BP1* KO cells were generated using a lentiviral-based CRISPR/Cas9 strategy, as previously described (Burigotto *et al*, 2021). Briefly, a gene-specific crRNA was cloned into the Lenti-CRISPR-V2 backbone (gift from Feng Zhang; Addgene plasmid #52961). This plasmid, along with pCMV-VSV-G (a gift from Bob Weinberg, Addgene plasmid #8454) and psPAX2 (a gift from Didier Trono, Addgene plasmid #12260), was used to co-transfect HEK 293T cells using calcium phosphate. After 48h, supernatants were harvested, filtered, mixed with 4 μg/mL hexadimethrine bromide (Sigma-Aldrich) and administered to cells for 24 h. Transduced cells were enriched by 10 μg/mL puromycin selection (InvivoGen) for 72 h. RPE1 PLK1^AS^ *TP53BP1* KO cells were generated using an RNP-based CRISPR/Cas9 strategy (Ghetti *et al*, 2021), as described above (without the addition of a single-strand DNA homology template nor NU-7741 treatment). For both cell lines, isogenic clones were isolated by limiting dilution. The presence of gene-disrupting INDELs in edited cells was confirmed by Sanger sequencing of PCR products spanning the crRNA recognition site, followed by Inference of CRISPR Edits (ICE) analysis (https://ice.synthego.com) (Hsiau *et al*, 2018).

### Generation of stable cell lines

To generate fluorescent histone H2B-labelled RPE1 p21-EGFP cells, pLentiPGK DEST H2B-iRFP670 (Addgene plasmid 90237) was introduced via lentiviral-mediated transduction. Cells with high transgene expression were isolated using FACS.

### Yeast two-hybrid screen

The yeast two-hybrid screen was performed by Hybrigenics Services, S.A.S., Evry, France. The KT binding domain sequence of 53BP1 (aa 1235-1616) was PCR-amplified from pcDNA5-FRT/TO-eGFP-53BP1 (Addgene plasmid #60813) and cloned into pB66 downstream to the Gal4 DNA-binding domain. This construct was used as bait to screen a human lung cancer cells cDNA library constructed into pP6. 70 million clones were screened using a mating approach with YHGX13 (Y187 ade2-101::loxP-kanMX-loxP, matα) and CG1945 (mata) yeast strains as previously described (Fromont-Racine *et al*, 1997). 173 His^+^ colonies were selected on a medium lacking tryptophan, leucine and histidine. Prey fragments of positive clones were amplified by PCR and sequenced at their 5’ and 3’ junctions. The resulting sequences were used to identify the corresponding interacting proteins in the GenBank database (NCBI) using a fully automated procedure. The sequences of 67 CENP-F fragments were overlapped and their positions calculated relative to the full-length protein sequence. The minimal region that overlaps represents the selected interacting domain.

### Clonogenic assay

4×10^3^ RPE1 cells were seeded in 10-cm dishes and medium was refreshed every 5 days. When applied, centrinone was refreshed every 3 days. After 8 - 16 days, cells were washed in PBS and then stained for 30 min at 37°C with crystal violet (0.2% w/v, AcrosOrganics) dissolved in 20% v/v methanol. Cell plates were rinsed thoroughly with water and left to dry overnight. Images were acquired using a ChemiDoc Imaging System (Bio-Rad).

### Cell lysis and Western blot

Cells were harvested by trypsinization and lysed in 50 mM Tris pH 7.4, 150 mM NaCl, 0.5% v/v NP-40, 50 mM NaF, 1 mM Na_3_VO_4_, 1 mM PMSF, one tablet/50 mL cOmplete, EDTA-free Protease Inhibitor Cocktail (Roche), 2 mM MgCl2 and 0.2 mg/mL DNase I (Thermo Fisher Scientific). Protein concentration was assessed by bicinchoninic acid assay (Pierce BCA Protein Assay Kit, Thermo Fisher Scientific) using a plate reader. Equal amounts of total protein samples (40-50 µg) were resolved by polyacrylamide gels using self-made or pre-cast gels (Bio-Rad). Proteins were electroblotted on nitrocellulose membranes (Cytiva) using a wet transfer system (Bio-Rad). Membranes were blocked in blocking solution (5% w/v non-fat milk in PBS-Tween 0.1% v/v), incubated overnight at 4°C with the relevant antibody diluted in blocking solution, washed several times with PBS-Tween 0.1% v/v and then incubated at room temperature for 50 min with HRP-conjugated secondary antibodies diluted in blocking solution. Protein detection was carried out using an Alliance LD2 Imaging System (UviTec Cambridge), after incubating membranes with Amersham ECL Select Western Blotting Detection Reagent (Cytiva). The following antibodies were used: rabbit anti-CENPF (Cell Signaling Technology, 58982,1:500), rabbit anti-53BP1 (Cell Signaling Technology, 4937, 1:500), mouse anti-cyclin A2 (Abcam, ab38, 1:500), mouse anti-vinculin (Sigma-Aldrich, V9264, 1: 2000), goat anti-rabbit IgG/HRP (Dako, P0448, 1:5000), rabbit anti-mouse IgG/HRP (Dako, P0161, 1:5000).

### Immunoprecipitation and MS analysis

WT and 53BP1-V5 tagged RPE1 cells were grown in 15-cm dishes, arrested in thymidine and released in nocodazole for 10 h. Mitotic shake-off was performed, cells were washed with PBS and lysed as described above. 700 µg of each sample were incubated with 30 µL of slurry Affi-Prep protein A resin (Bio-Rad) and 1 µg of mouse anti-V5 tag antibody (Invitrogen, R96025) for 4 h at 4°C. The beads were collected by centrifugation and washed twice with lysis buffer. Dried bead complexes were eluted by boiling samples in Bolt™ LDS sample buffer with 10% v/v Bolt™ Sample Reducing Agent (Thermo Fisher Scientific) at 95°C for 10 min. Samples were separated in precast 10% Bolt Bis-Tris Plus gel (Thermo Fisher Scientific), run for ∼1 cm and stained with a Coomassie solution. For each sample, the entire stained area was excised and washed with 100 mM NH_4_HCO_3_ in 50% v/v acetonitrile (ACN) for 15 min. The colourless gel plugs were then dehydrated by adding 100% ACN. The dried gel pieces were reduced with 10 mM DTT for 1 h and alkylated with 55 mM iodoacetamide for 30 min. Gel fragments were then washed in water, dehydrated with ACN, and incubated in a digestion solution containing 12.5 ng/µL trypsin (Thermo Fisher Scientific) in 100 mM NH_4_HCO_3_ at 37°C overnight. Peptides were extracted by sequentially treating gel pieces with 3% trifluoroacetic acid (TFA) in 30% v/v ACN, and 100% v/v ACN. Tryptic peptides were then dried in a speed-vac and acidified with TFA to a pH <2.5. After desalting on C18 stage tips, peptides were resuspended in 0.1% v/v formic acid (FA) for LC-MS/MS analysis.

Peptides were separated on an Easy-nLC 1200 HPLC system (Thermo Scientific) using a 25 cm reversed-phase column (inner diameter 75 µm packed in-house with ReproSil-Pur C18-AQ material: 3 µm particle size, Dr. Maisch, GmbH) with a two-component mobile phase (0.1% v/v FA in water and 0.1% v/v FA in ACN). Peptides were then eluted using a gradient of 5% to 25% over 50 minutes, followed by 25% to 40% over 15 min and 40% to 98% over 10 min at a flow rate of 400 nL/min. Peptides were analysed in an Orbitrap Fusion Tribrid mass spectrometer (Thermo Fisher Scientific) in data-dependent mode, with a full-scan performed at 120.000 FWHM resolving power (mass range: 350–1100 m/z, AGC target value: 10e6 ions, maximum injection time: 50 ms), followed by a set of (HCD) MS/MS scans over 3 sec cycle time at a collision energy of 30% (AGC target: 5 × 10e3 ions, maximum injection time: 150 ms). Dynamic exclusion was enabled and set at 30 sec, with a mass tolerance of 5 ppm. Data were acquired using Xcalibur 4.3 software and Tune 3.3 (Thermo Scientific). For all acquisitions, QCloud (Chiva *et al*, 2018) was used to control instrument longitudinal performance during the project using in-house quality control standards. Raw files were searched using Proteome Discoverer v.2.2.0 (Thermo Scientific). Peptide searches were performed against the in-silico digested UniProt Human database (downloaded April 2021), added with major known contaminants and the reversed versions of each sequence. Trypsin/P was chosen as the enzyme with 5 missed cleavages, and static modification of carbamidomethyl (C) with variable modification of oxidation (M) and acetylation of protein N-terminus were incorporated in the search. The MASCOT search engine (v.2.6.2, MatrixScience) was used to identify proteins (precursor mass tolerance: 10 ppm, product mass tolerance: 0.6 Da). The FDR was set to < 0.01 at both the peptide and protein levels. Peak intensities of the peptides were log_2_ transformed and data were normalised on the average of the specific protein abundance within each sample (Aguilan *et al*, 2020). The fold change (FC) of each peptide was calculated. Then, the FC at protein level was calculated by averaging the FC of all peptides assigned to each protein. 53BP1-binding partners were identified by subtracting the log_2_-normalised intensities of the control sample (RPE1 WT mitotic cells) to the test sample (RPE1 53BP1-V5 mitotic cells). Statistical significance was assessed using Student’s t-test (two-tailed, two-sample unequal variance).

### Competition assay and FACS analysis

To generate cells constitutively expressing EGFP, the EGFP sequence was PCR amplified from pEGFP-N1 (Clontech), and cloned in the pAIB-CAG lentiviral vector (a gift from Claudio Ballabio) between the PmeI and NotI sites, upstream of an IRES element and a blasticidin resistance gene under the CAG promoter. Lentiviral particles were produced as described above. After transduction, cells were treated with 5 μg/mL blasticidin for 7 days. For competition growth assays, RPE1 WT cells stably expressing EGFP and non-fluorescent RPE1 cells of the desired genotype were mixed at a 1:1 ratio and seeded into duplicate wells. One well from each pair was treated with centrinone for 8 days. Cells were analysed on a Symphony A1 cytometer (BD Biosciences) to assess the percentage of EGFP+ cells. Live cells were gated through forward-scatter (FSC) and side scatter (SSC) parameters, and cell doublets were excluded. For each sample, the fraction of EGFP-cells was divided by the fraction of EGFP+ cells. The value obtained from the centrinone treated well was then divided by that obtained in the untreated well to determine the fold change in EGFP-cells. Analysis of flow cytometry data was performed using FlowJo software (FlowJo, LLC).

### Immunofluorescence

Cells were grown on glass coverslips (Marienfeld-Superior), washed in PBS and directly fixed and permeabilized with absolute ice-cold methanol for at least 20 min at −20°C. For NudE staining cells were pre-extracted for 2 min in PTEM buffer (0.2% v/v Triton X-100, 20 mM PIPES at pH 6.8, 1 mM MgCl_2_, 10 mM EGTA in ddH_2_O) and then fixed with 4% v/v formaldehyde (Sigma-Aldrich) in PTEM for 10 min at room temperature. Cells were rinsed with PBS, blocked with 3% w/v BSA in PBS for 20 min and stained for 1 h at room temperature with primary antibodies diluted in blocking solution. Cells were washed with PBS and incubated with fluorescent secondary antibodies for 45 min at room temperature. DNA was stained with 1 µg/mL Hoechst 33342 (Invitrogen). After incubation, cells were rinsed with PBS and ddH2O, and mounted using ProLong Gold Antifade Reagent (Invitrogen). The following antibodies were used: mouse anti-53BP1 (Millipore, MAB3802, 1:500), mouse anti-V5 (Invitrogen, R96025, 1:1000), rabbit anti-CENPF (Cell Signaling Technology, 58982,1:500), human anti-centromere/KT (Antibodies Inc, 15-234, 1:500), rabbit anti-NudE (ProteinTech, 10233-1-AP, 1: 250), mouse anti-phospho-Histone H2A.X Ser139 (Millipore, 05-636, 1:500), goat anti-mouse AlexaFluor488 (Invitrogen, A11029, 1:1000), goat anti-mouse AlexaFluor 555 (Invitrogen, A21424, 1:1000), goat anti-rabbit AlexaFluor 488 (Invitrogen, A11034, 1:1000), goat anti-rabbit AlexaFluor 555 (Invitrogen, A21429, 1:1000), goat anti-human AlexaFluor 647 (Invitrogen, A21445, 1:1000). For Fig. 1H the following antibodies were used: sheep anti-CENP-F (kind gift of S. Taylor, University of Manchester, SCF.1, 1:1000), mouse anti-CENP-A (GeneTex, GTX13939, 1:2000), rabbit anti-53BP1 (Novus Biologicals, NB100-304, 1:500), donkey anti-rabbit AlexaFluor 488 (Thermo Fisher, A21206, 1:1000), donkey anti-mouse AlexaFluor 555 (Thermo Fisher, A31570, 1:1000), donkey anti-sheep AlexaFluor 647 (Thermo Fisher, A21448, 1:1000). Images were acquired on a spinning disc Eclipse Ti2 inverted microscope (Nikon Instruments Inc), equipped with Lumencor Spectra X Illuminator as LED light source, an X-Light V2 Confocal Imager and an Andor Zyla 4.2 PLUS sCMOS monochromatic camera using a plan apochromatic 100×/1.45 oil immersion objective. Images in Fig. 1H were collected using a Deltavision Elite system (GE Healthcare) controlling a Scientific CMOS camera (pco.edge 5.5).

Acquisition parameters were controlled by SoftWoRx suite (GE Healthcare). Images were collected using an Olympus 100×/1.4 oil immersion objective. Images in Fig. EV1A, and Appendix Fig. A and C were acquired on a Nikon AX confocal microscope (Nikon Instruments Inc) equipped with a LUA-S4 laser unit using a plan apochromatic 100×/1.49 oil immersion objective. Images were deconvolved with Huygens Professional software (Scientific Volume Imaging, Hilversum, The Netherlands). Images were processed using Fiji and displayed as maximum intensity projections of deconvolved z-stacks. All displayed images were selected to most closely represent the mean quantified data.

### Mitotic Surveillance Pathway threshold assay

RPE1 p21-EGFP H2B-iRFP cells were treated with a siRNA SMART pool targeting CENP-F or a non-targeting control for 24 hr. Cells were then seeded into 4-well chamber slides (Ibidi) for a further 24 hr. Before imaging, standard media was replaced with CO_2_-independent base medium (Thermo Fischer) and maintained at 37 °C in an environmental control station. Once under the microscope, cells were blocked in mitosis with 2 µM dimethylenastron for 6.5 h. Fresh media was then replaced to allow cells to exit mitosis. Daughter cell senescence was monitored for 2 days using p21-EGFP expression and related to time spend in mitosis (NEBD-anaphase onset). Long-term time-lapse imaging was performed using a Deltavision Elite system (GE Healthcare) controlling a Scientific CMOS camera (pco.edge 5.5). Images were acquired with an Olympus 40× 1.4 NA oil objective.

### Image analysis and statistics

All images of similarly stained experiments were acquired with identical illumination settings. To quantify fluorescence intensity at KTs, maximum intensity projections of z-stacks were obtained using Fiji and circular regions of interest (ROIs) with a diameter of 5 pixels were drawn, centered on the KT (CREST signal). Measurements were performed on at least 10 independent images. For each cell, 3 background ROIs (placed in the cytoplasmic space, in proximity to the DNA staining) were drawn and their average value was subtracted from the intensity value of each KT. To quantify DNA damage foci, cell nuclei were segmented using Fiji, according to the Hoechst signal. The “find maxima” function was then applied to quantify the number of foci inside each nucleus. To quantify the number of colonies in the clonogenic assay, plates scans were imported in Fiji, manually cropped to individual plates and median filtered. Colonies were equally segmented among different genotypes and treatments, and the “analyse particles” function was then used to count colonies. Data are presented as dot plot (mean ± SD). Graphs were produced using GraphPad Prism 9 (GraphPad, San Diego, CA, USA) software.

## Acknowledgements

We thank all members of the Fava laboratory for useful discussions and Dr. Matthias Altmeyer for suggestions. We also thank Drs. Giorgina Scarduelli, Michela Roccuzzo, Daniele Peroni, and Romina Belli for technical support. The research leading to these results has received funding from the Giovanni Armenise-Harvard Foundation (CDA 2017), from the University of Trento, from the MIUR PRIN 2017 (CUP E64I19001070001), from AIRC (under MFAG 2019, project ID. 23560) and from Telethon (GJC21181), P.I. Luca Fava.

## Expanded View Figure legends

**Figure EV1.**
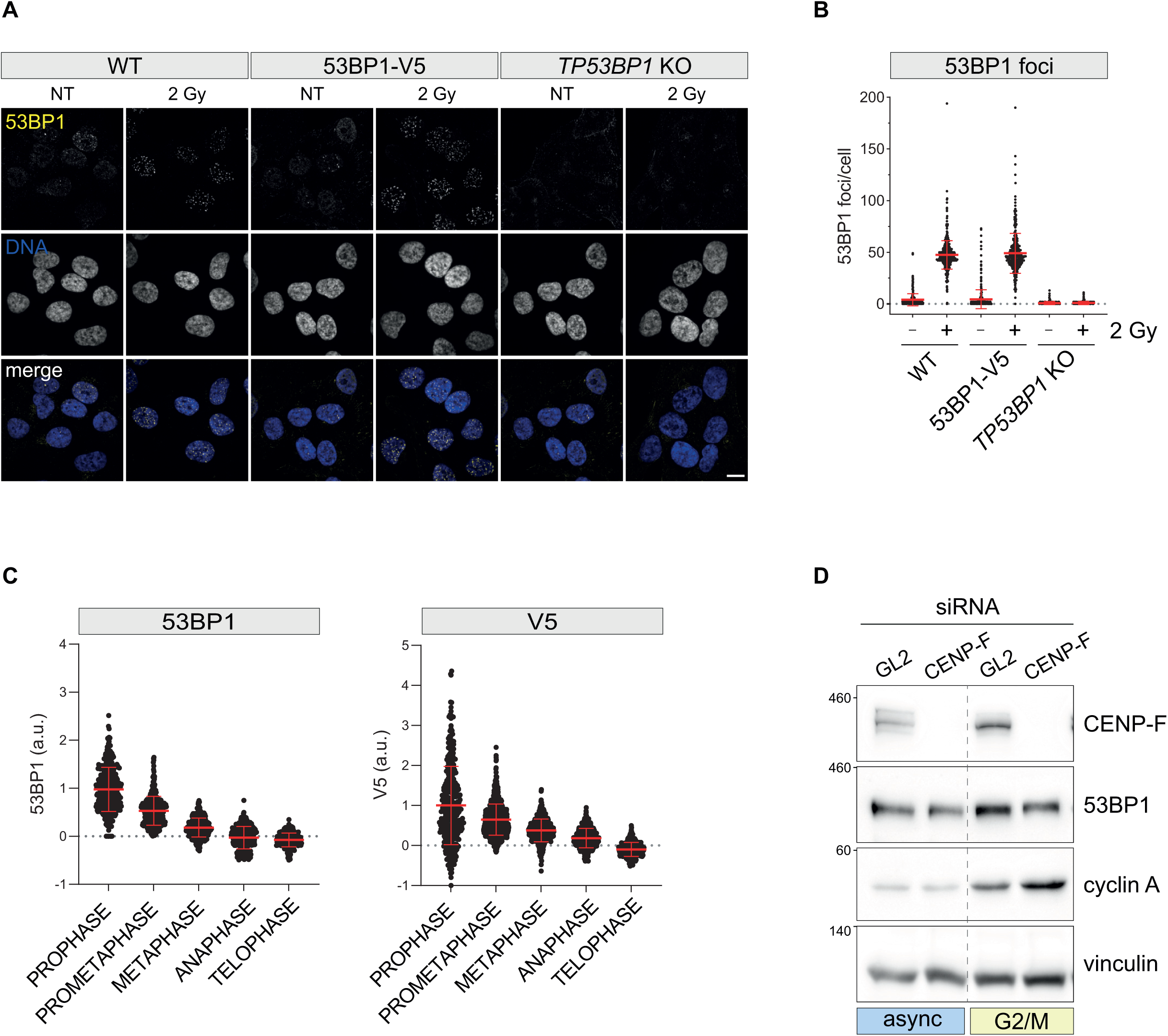
A) Representative fluorescence micrographs of cells subjected to irradiation (2 Gy) or left untreated and probed against 53BP1 protein. Scale bar: 10 µm. B) Dot plots showing the number of 53BP1 foci in cells treated as in A). Mean values (red lines) ± SD are reported. N ≥ 352 cells were quantified for each condition. C) Dot plots showing 53BP1 (in RPE1 WT cells, left panel) or V5 (in RPE1 53BP1-V5 cells, right panel) average pixel intensity at individual KTs across the indicated cell cycle phases. Mean values (red lines) ± SD are reported. N ≥ 349 KTs were assessed from 10 cells for each mitotic phase; a.u. = arbitrary units. D) HeLa S3 cells were transfected with the indicated siRNA and either treated with thymidine for 24 h and released in fresh medium for 10 h (G_2_/M), or left untreated (async = asynchronous). Cells were subjected to immunoblotting using the indicated antibodies.

**Figure EV2.**
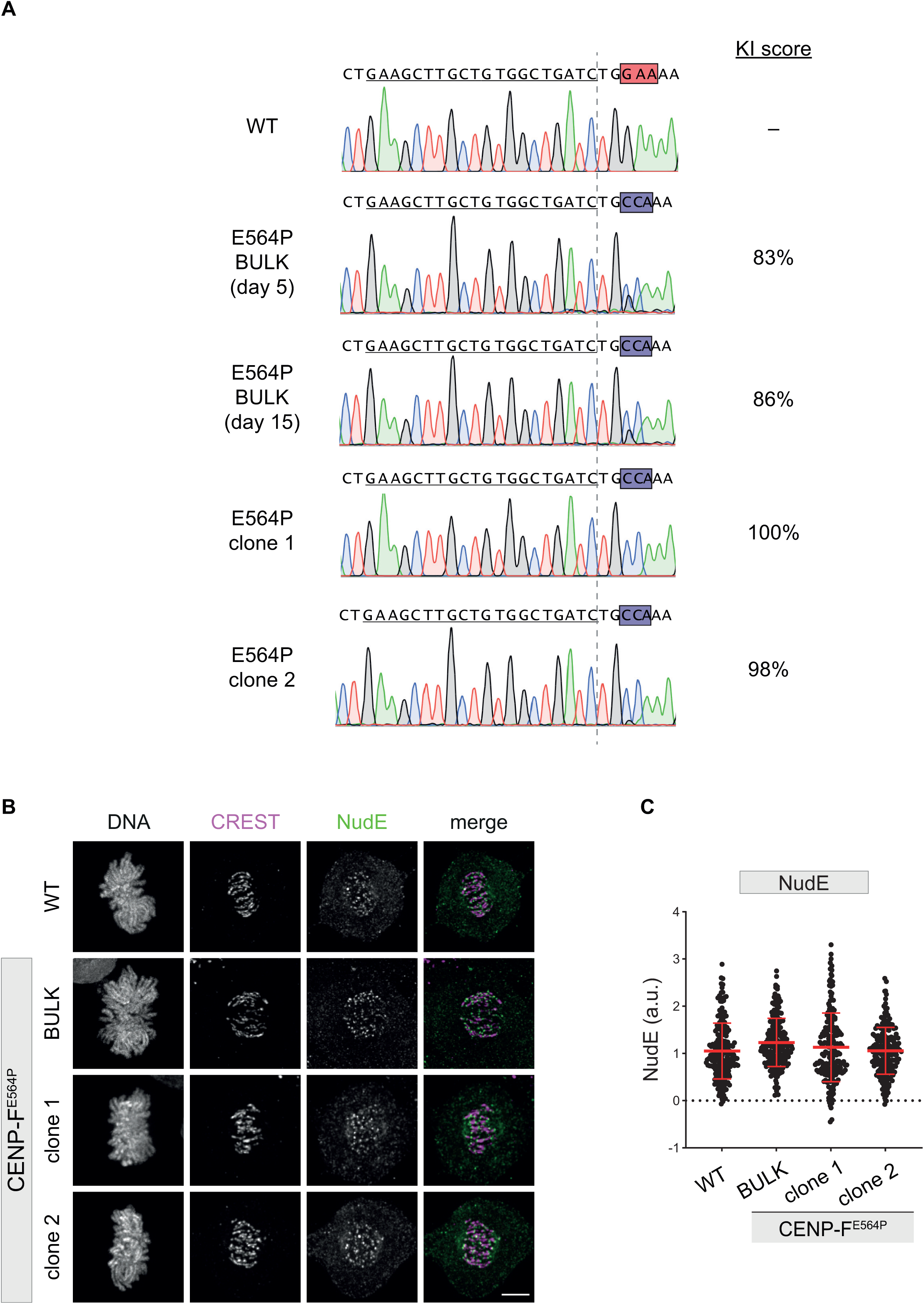
A) The *CENPF* region targeted by the gRNA (black solid line) was PCR amplified and sequenced from genomic DNA of the indicated cells. Electropherograms are shown, along with the knock-in score calculated by ICE. Vertical dashed line: Cas9 cut site; red box: WT codon (E564); blue box: mutant codon (P564). B) Fluorescent micrographs of RPE1 cells of the indicated genotype, co-stained with the indicated antibodies. Scale bar: 5 µm. C) Dot plots showing the intensity of NudE protein at individual KTs. Mean values (red lines) ± SD are reported. Data obtained from images as in B). N ≥ 211 KTs were assessed from at least 10 cells for each condition; a.u. = arbitrary units.

**Figure EV3.**
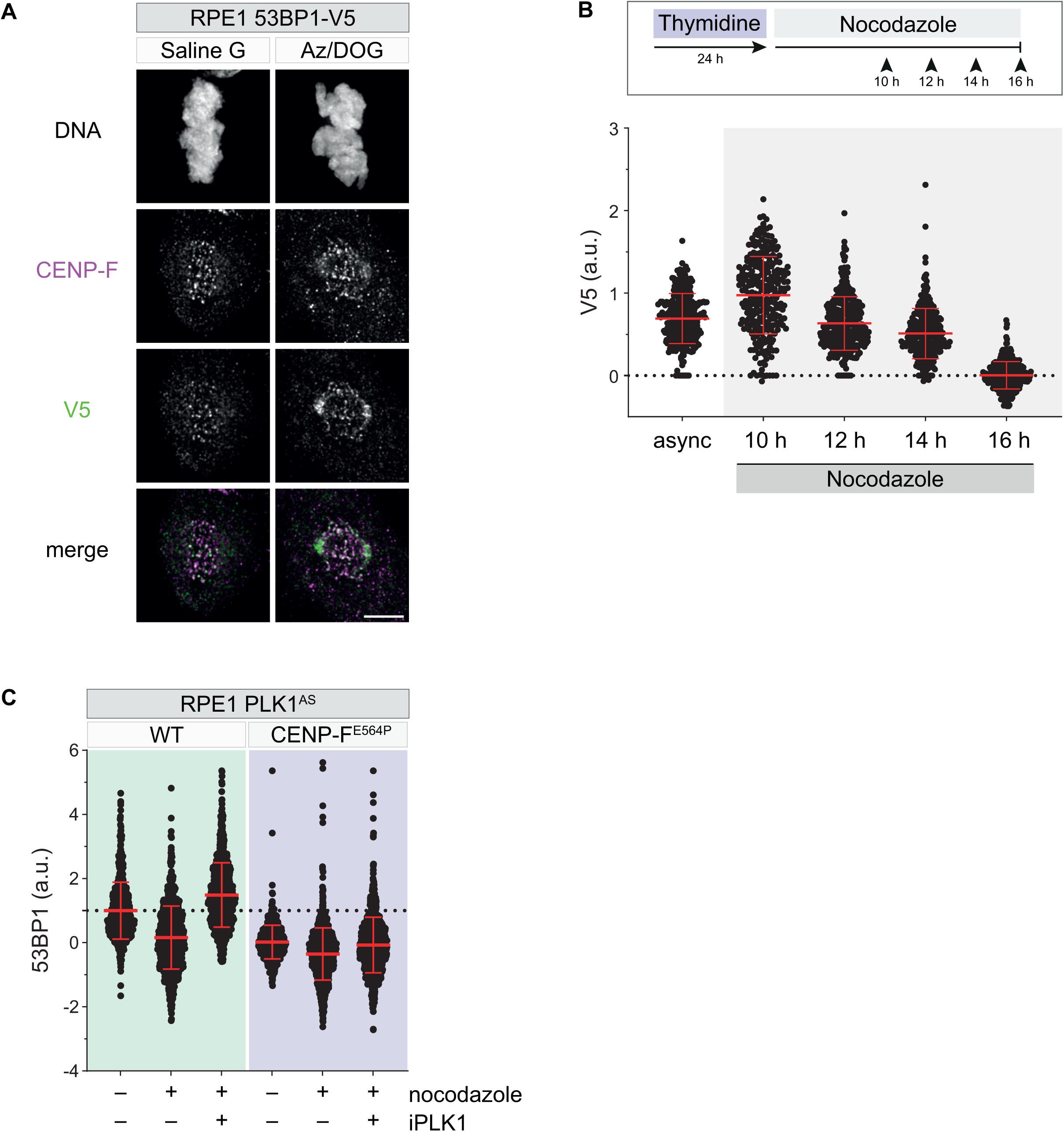
A) Representative fluorescence micrographs of RPE1 53BP1-V5 cells incubated for 15 min in isotonic salt solution in the presence of either glucose (Saline G) or sodium azide/2-deoxy-D-glucose (AZ/DOG). Scale bar: 5 µm. B) Top: RPE1 53BP1-V5 cells were treated with thymidine for 24 h, released in medium containing 3.3 µM nocodazole and fixed after 10-16 h. Bottom: dot plots showing V5 fluorescence intensity at individual KTs. Mean values (red lines) ± SD are reported. N ≥ 266 KTs were assessed from at least 20 cells for each condition; async = asynchronous cells; a.u. = arbitrary units. C) Dot plots showing the intensity of 53BP1 fluorescence at individual KTs. Mean values (red lines) ± SD are reported. Data obtained from images as in Figure 4C. N ≥ 849 KTs were assessed from at least 20 cells for each condition; a.u. = arbitrary units.

## Appendix Figure Legends

A) Representative fluorescence micrographs of cells subjected to irradiation (2 Gy) or left untreated and probed against γH2AX protein. Scale bar: 10 µm. B) Dot plots showing the number of γH2AX foci in cells treated as in A). Mean values (red lines) ± SD are reported. N ≥ 280 cells were quantified for each condition. C) Representative fluorescence micrographs of cells treated as in A) and probed against V5-tag. Scale bar: 10 µm. D) Dot plots showing the number of V5 foci in cells treated as in A). Mean values (red lines) ± SD are reported. N ≥ 349 cells were quantified for each condition.

